# Ire1 membrane stress responses support cell growth upon disruptions in inter-organelle contacts

**DOI:** 10.64898/2026.07.27.740907

**Authors:** Bailey R.W. Hewlett, Bianca M. Esch, Ffion B. Thomas, Misako Araki, Atsuko Ikeda, Kazuki Hanaoka, Kouichi Funato, Florian Fröhlich, Christopher J. Stefan

**Affiliations:** Laboratory for Molecular Cell Biology, University College London, Gower Street, London WC1E 6BT, UK; Department of Biology/Chemistry, Bioanalytical Chemistry Section, University of Osnabrück, Osnabrück, Germany; Center for Cellular Nanoanalytic Osnabrück (CellNanOs), Osnabrück University, Germany; Graduate School of Integrated Sciences for Life, Hiroshima University, Kagamiyama 1-4-4, Higashi-Hiroshima 739-8528, Japan

**Author notes:** **Correspondence:** Christopher J. Stefan. Equal contributions (Bailey R.W. Hewlett, Bianca M. Esch).

**Keywords:** inter-organelle contacts, lipid transfer proteins, membrane stress responses, phospholipid homeostasis, sphingolipid homeostasis

## Abstract

Disruptions in inter-organelle contacts result in membrane lipid homeostasis defects that ultimately impair cellular function and viability. Yet, essential responses to membrane lipid imbalances remain poorly understood. In this study, we demonstrate that Ire1-dependent Membrane Stress Responses (MSR), distinct from the canonical Unfolded Protein Response (UPR), sustain the growth of yeast cells lacking inter-organelle contacts. Comprehensive lipidomics reveal that the Ire1-mediated MSR compensates for glycerophospholipid synthesis defects by modulating sphingolipid metabolism. Quantitative imaging further indicates that Ire1 mediates these effects, at least in part, by elevating cytoplasmic Ca^2+^ which in turn stimulates calcineurin activity necessary for cellular homeostasis. Accordingly, inhibition of calcineurin results in severe endoplasmic reticulum stress in yeast cells depleted of inter-organelle contacts. Thus, the Ire1 MSR directs Ca^2+^-dependent calcineurin activity and lipid metabolism upon disruptions in membrane contact sites to maintain cellular homeostasis. Alterations in inter-organelle contacts are associated with several diseases, including neurodegenerative disorders. Our findings in yeast suggest that evoking the MSR may be a means to ameliorate neuronal degeneration and delay the progression of neurodegenerative disorders.

## Introduction

Inter-organelle contacts between the endoplasmic reticulum (ER) and other membrane-bounded compartments, including mitochondria, endosomes, lysosomes, and Golgi compartments, are important sites for non-vesicular lipid and calcium (Ca^2+^) transport and homeostasis (Nishimura and Stefan, 2020, Prinz et al., 2020, Voeltz et al., 2024). Consequently, disruptions in non-vesicular transport at inter-organelle contacts result in severe mitochondrial, endo-lysosomal, and secretory dysfunction. However, less is known about how ER function may be impacted, or whether and how ER stress responses might restore homeostasis upon alterations in membrane contact sites.

In the budding yeast *Saccharomyces cerevisiae*, ER stress responses are primarily mediated by the highly conserved transmembrane protein Ire1 (IRE1α ortholog). Misfolded proteins in the ER engage with Kar2 (BiP) that attenuates Ire1 (Okamura et al., 2000) and directly bind the N-terminal luminal domain of Ire1 (Gardner and Walter, 2011) promoting Ire1 dimerization and oligomerization that induce conformational changes in the Ire1 cytoplasmic kinase and site-specific endoribonuclease (RNase) domains (Aragon et al., 2009, Korennykh et al., 2009). Ire1-dependent *HAC1* mRNA splicing subsequently results in expression of the Hac1 transcription factor (XBP1 ortholog), which elicits a transcriptional response termed the Unfolded Protein Response (UPR) (Cox and Walter, 1996, Travers et al., 2000, Walter and Ron, 2011). In addition, Ire1 and IRE1α can detect changes in the lipid composition and biophysical properties of the ER membrane bilayer, such as lipid acyl chain saturation, via their transmembrane and juxtamembrane regions (Ariyama et al., 2010, Promlek et al., 2011, Volmer et al., 2013, Halbleib et al., 2017, Kono et al., 2017, Ho et al., 2020). However, Ire1 and IRE1α dimers do not cluster into higher-order oligomers in response to membrane lipid stress (Ishiwata-Kimata et al., 2013, Kitai et al., 2013, Halbleib et al., 2017, Ho et al., 2020), resulting in a more moderate level of Ire1 activity that reportedly elicits a distinct transcriptional program termed the Membrane Stress Response (MSR) (Thibault et al., 2012, Fun and Thibault, 2020, Ho et al., 2020). Although previous studies have suggested that Ire1-mediated MSR does not restore membrane bilayer homeostasis upon disruptions in phospholipid synthesis (Thibault et al., 2012, Ho et al., 2020), Ire1 has been implicated in the control of phospholipid synthesis and acyl chain saturation in the ER membrane bilayer (Travers et al., 2000, Schuck et al., 2009, Surma et al., 2013, Volmer and Ron, 2015). Thus, the full extent and roles of the Ire1-mediated MSR are not completely understood.

In addition to elucidation of the Ire1-dependent UPR and MSR pathways, studies in budding yeast have provided instrumental insights into the roles of membrane contact sites in the control of lipid and Ca^2+^ homeostasis. For example, several studies have shown that loss of the yeast Scs2 protein (VAP ortholog) and its paralog Scs22 along with several additional inter-organelle contact site proteins including Ist2 (ortholog of the ANO8/TMEM16H and ANO10/TMEM16K Ca^2+^-activated lipid scramblases) and the tricalbin proteins (Tcb1/2/3, orthologs of the extended synaptotagmin E-Syt1/2/3 Ca^2+^-activated lipid transfer proteins) results in disconnection of the cortical ER network from the plasma membrane (PM) as well as from other organelles (Loewen et al., 2007, Manford et al., 2012, Quon et al., 2018, Collado et al., 2019, Hoffmann et al., 2019). These disruptions lead to alterations in membrane lipid metabolism and distribution, as well as elevations in cytoplasmic Ca^2+^ signals (Loewen et al., 2007, Manford et al., 2012, Omnus et al., 2016, Kato et al., 2017, Quon et al., 2018, Nishimura et al., 2019, Jorgensen et al., 2020, Ikeda et al., 2020, Quon et al., 2022, Thomas et al., 2022, Nenadic et al., 2023, Hanaoka et al., 2024). One previous study in particular revealed that cells lacking six ER-localized membrane contact site proteins (Scs2/22, Ist2, and Tcb1/2/3), named Δtether cells to emphasize the drastic reductions in ER-PM contacts, resulted in constitutive Ire1-dependent ER stress responses necessary for normal cell growth (Manford et al., 2012). However, the causes of ER stress (proteotoxicity, membrane lipid imbalances, or some other trigger) were not addressed., Furthermore, it was unclear whether the UPR or the MSR (or both) is the Ire1-dependent response required for growth of the Δtether cells.

In the present study, we address these issues and demonstrate, first, that robust UPR is not essential in the Δtether yeast cells, indicating that the moderate Ire1 MSR is sufficient for growth of the Δtether cells. Second, using quantitative lipidomics, we show that the Δtether cells have significantly reduced levels of mono-unsaturated phosphatidylserine and phosphatidylethanolamine species, resulting in a shift in the acyl chain saturation index of these glycerophospholipids to di-unsaturated species. These effects are exacerbated upon loss of Ire1 in the Δtether cells and rescued by the Ire1 MSR. Moreover, concomitant with the reduction in mono-unsaturated glycerophospholipids, the Ire1 MSR is necessary and sufficient for markedly increased production of the saturated long chain sphingoid bases dihydrosphingosine and phytosphingosine, which are normally converted to ceramides and more complex sphingolipids, but can also be utilized as a source of saturated fatty acids for glycerophospholipid synthesis. This Ire1 MSR-dependent change in sphingolipid metabolism in the ER is due to an increase in Ca^2+^-mediated stimulation of calcineurin, which results in a decrease in ceramide synthase function due to dephosphorylation of its catalytic subunits Lac1 and Lag1 (Aronova et al., 2008, Muir et al., 2014, Omnus et al., 2016). Consistent with this conclusion, we show that interference of this homeostatic regulatory loop with the calcineurin inhibitor FK506 results in severe ER stress in the yeast Δtether cells. Thus, upon disruptions in membrane contact sites, the Ire1 MSR redirects lipid metabolic channelling, from impaired glycerophospholipid production to increased sphingoid base synthesis in the ER, in a manner that circumvents ceramide toxicity and chronic ER stress that may lead to cell death if unresolved.

## Results

### Ire1 membrane stress responses (MSR) are sufficient for cell growth upon disruptions in inter-organelle contacts

In budding yeast, loss of Scs2/22 (VAP-A/B orthologs), Ist2 (ANO8/ANO10 ortholog), and the Tcb1/2/3 proteins (E-SYT1/2/3 orthologs) markedly decreases ER-PM contact formation and causes alterations in membrane lipid and Ca^2+^ homeostasis (Loewen et al., 2007, Wolf et al., 2012, Toulmay and Prinz, 2012, Manford et al., 2012, Omnus et al., 2016, Obara and Kihara, 2017, Kato et al., 2017, Quon et al., 2018, Nishimura et al., 2019, D’Ambrosio et al., 2020, Jorgensen et al., 2020, Ikeda et al., 2020, Wong et al., 2021, Thomas et al., 2022, Nenadic et al., 2023, Hanaoka et al., 2024), resulting in prominent defects in PM and ER homeostasis (Figure 1A). In particular, a previous study (Manford et al., 2012) showed that cells lacking the Scs2/22, Ist2, and Tcb1/2/3 proteins, named Δtether cells, display significant changes in ER morphology as well as elevated Ire1-dependent ER stress responses, even under normal growth conditions (see Figure 1B). Moreover, it has been reported previously that loss of Ire1 or the Ire1-regulated Hac1 transcription factor severely impairs growth of the Δtether cells (Manford et al., 2012, Jorgensen et al., 2020). Ire1 elicits the MSR in response to membrane stresses and the UPR in response to protein misfolding in the ER lumen (see Figure 1D). Yet, to date, it has been unclear what leads to constitutive Ire1 activity in the Δtether cells and how Ire1-dependent ER stress responses support growth of the Δtether cells.

**Figure 1.**
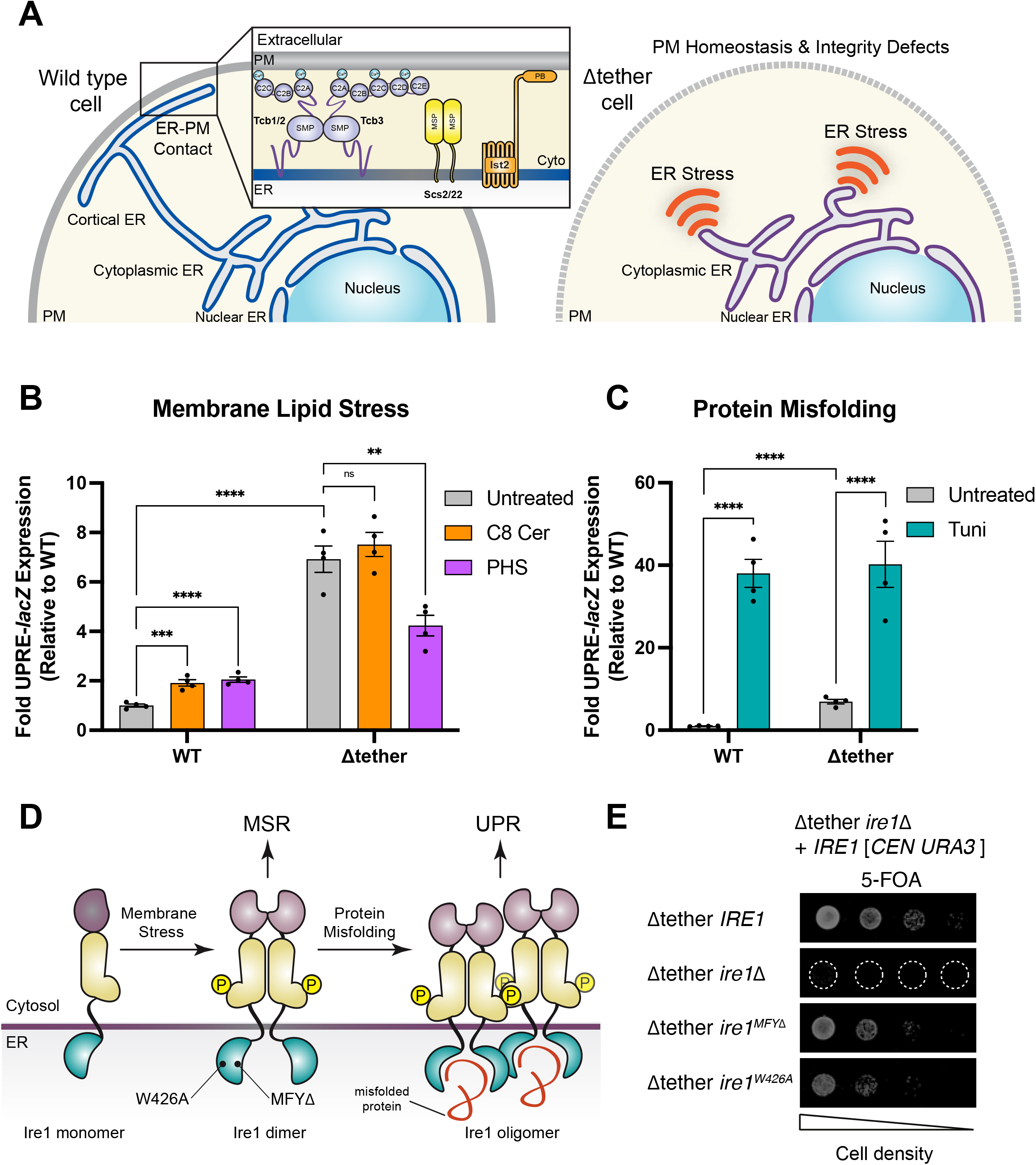
Ire1-mediated membrane stress responses (MSR) are upregulated and sufficient for cell growth upon loss of ER-PM contacts. (A) Cartoon representation of proteins localised to ER-PM contacts in budding yeast. The Δtether mutant cells lack the tricalbins (Tcb1/2/3), Scs2/22 and Ist2 proteins, causing the ER to detach from the PM and other organelles resulting in membrane homeostasis defects and ER stress. (B) Ire1 activity was monitored by expression of a *UPRE-lacZ* reporter in wild-type and Δtether cells treated with C8-Ceramide (C8-Cer, 100 μg/ml) or PHS (50 μg/ml) to induce membrane stress responses (MSR). Data represent mean +/-standard error (N=4 independent experiments). Standard deviation is provided in the Data Table. (C) Tunicamycin (Tuni, 2 μg/ml) induces protein misfolding and robust unfolded protein responses (UPR) in wild-type and Δtether cells, as monitored by expression of a *UPRE-lacZ* reporter. Data represent mean +/-standard error (N=4 independent experiments). Standard deviation is provided in the Data Table. (D) Cartoon representation of Ire1 dimerization in response to membrane lipid stress and oligomerisation in response to proteotoxic stress, respectively. Ire1 oligomers induce robust UPR upon accumulation of misfolded proteins in the ER, while oligomers are not formed during MSR. The Ire1^MFYΔ^ mutant is impaired in binding misfolded proteins in the ER; the Ire1^W426A^ mutant is impaired in forming oligomers but still can form dimers. (E) The Δtether *ire1*Δ cells exhibit a severe growth defect. Plasmid shuffle assays with serial dilutions (10-fold) of Δtether *ire1*Δ cells co-transformed with an *URA3*-marked *IRE1* plasmid and either an empty *LEU2*-marked *CEN* vector or vector expressing wild-type Ire1, Ire1MFYΔ, or Ire1W426A were spotted onto agar media containing 5-FOA to select against the *URA3*-marked *IRE1* plasmid. Expression of Ire1^MFYΔ^ or Ire1^W426A^, which are impaired in UPR, is sufficient to rescue the impaired growth of the Δtether *ire1*Δ cells. ****, p < 0.0001; ***, p < 0.0002; ** p < 0.0021; ns, not significant.

First, to assess the nature and extent of ER stress in the Δtether cells, we used a well-established *UPRE*-*lacZ* reporter assay (Mori et al., 1992, Cox and Walter, 1996, Mori et al., 1996) which monitors Ire1 and Hac1 activity. As a measure of the Ire1-mediated MSR, we challenged cells with two membrane perturbants, either short acyl chain ceramide (diC8 Cer) or the long chain base phytosphingosine (PHS), which induced a modest, but statistically significant, increase in *UPRE-lacZ* expression in wild-type cells (2-fold above that in growth medium alone, Figure 1B). Even in growth medium alone, the Δtether cells displayed a greater than 6-fold increase in basal *UPRE-lacZ* expression (Figure 1B), consistent with a previous study (Manford et al., 2012). However, in contrast to wild-type cells, the Δtether cells did not display further increases in *UPRE-lacZ* expression in response to diC8 Cer, and *UPRE-lacZ* expression was intriguingly reduced in Δtether cells exposed to PHS (Figure 1B; see Discussion). As a measure of the Ire1-mediated UPR, we exposed cells to the N-glycosylation inhibitor tunicamycin, which causes protein misfolding in the ER lumen (Kuo and Lampen, 1974). *UPRE-lacZ* expression increased significantly, but equivalently, in both wild-type and Δtether cells upon tunicamycin-induced protein misfolding in the ER (approximately 40-fold above wild-type control levels; Figure 1C). Altogether, these results suggested that the MSR is constitutively and maximally induced in the Δtether cells, whereas there is modest induction of the UPR.

To determine whether the Ire1-mediated MSR is sufficient to support growth of the Δtether cells, we took advantage of two Ire1 mutants shown to cripple the ability of Ire1 to elicit an effective UPR in response to protein misfolding: Ire1^MFYΔ^ that is impaired in binding misfolded proteins and Ire1^W426A^ that is impaired in the formation of higher-order oligomers (Gardner and Walter, 2011) (Figure 1D). The MSR does not rely on the Ire1 luminal domain or higher-order oligomerization. Instead, the MSR is primarily elicited by the transmembrane and juxtamembrane regions of Ire1 that mediate its dimerization upon perturbation of the ER membrane lipid environment (Promlek et al., 2011, Ishiwata-Kimata et al., 2013, Kitai et al., 2013, Volmer et al., 2013, Halbleib et al., 2017, Ho et al., 2020) (Figure 1D). We tested the capacity of Ire1^MFYΔ^ and Ire1^W426A^ to support the growth of the Δtether cells when present as the sole source of Ire1 using a plasmid shuffle approach. As a control, complete loss of Ire1 significantly impaired growth of the Δtether cells (Δtether *ire1*Δ; Figures 1E and S1A); the Δtether *ire1*Δ cells are still viable but grow very slowly, as previously reported (Manford et al., 2012). In contrast, even though Ire1^MFYΔ^ or Ire1^W426A^ are impaired in eliciting an effective UPR, they were able to support the growth of the Δtether cells nearly as well as wild-type Ire1 (Figures 1D and S1A). Furthermore, it is noteworthy that Ire1^MFYΔ^ or Ire1^W426A^ were able to support the growth of the Δtether cells, even though constitutive *UPRE-lacZ* expression increased by only 2-fold above wild-type basal levels in the Δtether *ire1MFY*Δ and Δtether *ire1W426A* cells (Figure S1B). Thus, importantly, these results revealed that only a modest Ire1-dependent stress response consistent with the MSR, and not a full-blown UPR, is sufficient to support the growth of cells lacking the membrane contact site proteins Scs2/22, Ist2, and Tcb1/2/3.

### Perturbations of the cellular lipidome in the Δtether mutant cells

Ire1-mediated UPR upregulates protein quality control and lipid biosynthesis pathways in response to proteotoxic stress in the ER (Travers et al., 2000, Schuck et al., 2009, Epstein et al., 2012). In contrast, Ire1-mediated MSR reportedly does not restore membrane lipid homeostasis upon disruptions in glycerophospholipid metabolism and instead is proposed to only upregulate protein quality control systems to maintain cellular viability (Thibault et al., 2012, Fun and Thibault, 2020, Ho et al., 2020). However, it is not known whether Ire1 MSR regulates membrane lipid homeostasis upon disruptions in inter-organelle contact sites that impair non-vesicular lipid transport from the ER.

To address whether the Ire1 MSR regulates lipid homeostasis in the Δtether cells, we determined steady-state levels (pmol lipid/μg protein) of diacylglycerol (DAG), triacylglycerol (TAG), and major glycerophospholipids (GPL) in wild-type, Δtether, Δtether *ire1*Δ, and Δtether *ire1W426A* cells by mass spectrometry-based lipidomics (Figures 2 and S2). Budding yeast cells can use two distinct pathways for *de novo* GPL synthesis (Henry et al., 2012), namely the cytidine diphosphate DAG (CDP-DAG) pathway and the Kennedy pathway (Figure 2A). Compared to wild-type cells, the Δtether cells displayed significant reductions in steady-state levels of the GPLs phosphatidylserine (PS), which is generated by the CDP-DAG pathway in yeast, as well as phosphatidylethanolamine (PE) and phosphatidylcholine (PC), which are generated by both the CDP-DAG and Kennedy pathways (Figure 2B), similar to previous reports (Quon et al., 2018, Nishimura et al., 2019, Jorgensen et al., 2020, Thomas et al., 2022). Disruptions in GPL synthesis can result in potentially toxic accumulation of DAG, if it is not converted into TAG (Rockenfeller and Gourlay, 2018). Previous studies have reported an increase in DAG levels in the Δtether cells (Quon et al., 2018, Jorgensen et al., 2020), suggesting that the Ire1 MSR may prevent DAG lipotoxicity. However, as compared to wild-type cells, we observed a slight decrease in DAG levels in the Δtether cells and no significant increase in DAG levels in the Δtether *ire1*Δ cells (Figure 2B). In addition, there were modest elevations in TAG levels in the Δtether, Δtether *ire1*Δ, and Δtether *ire1W426A* cells, as compared to wild-type cells (Figure 2B). Thus, the Ire1 MSR is not required to alleviate accumulation of a toxic level of DAG and is not required to produce TAG in the Δtether cells (Figure 2B).

**Figure 2.**
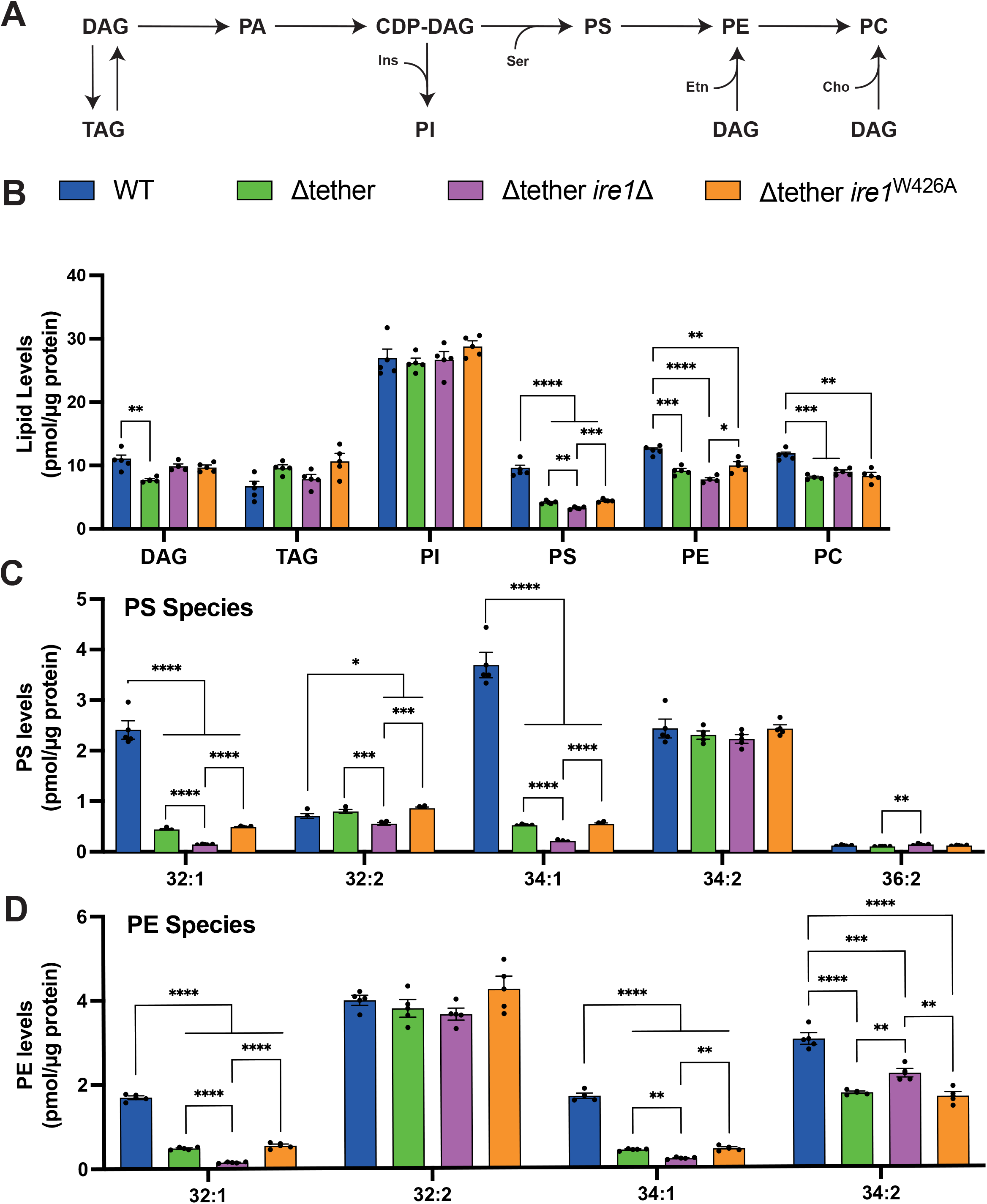
Ire1-mediated membrane stress responses (MSR) sustain pools of glycerophospholipids upon loss of ER-PM contacts. (A) Schematic of major glycerophospholipid (GPL) biosynthesis pathways in budding yeast. (B) Steady-state levels of diacylglyerol, triacylglyerol, and major GPLs in wild-type (WT), Δtether, Δtether *ire1*Δ and Δtether *ire1*^W426A^ cells as determined by quantitative mass spectrometry. (C and D) Species-level lipidomics analysis (total acyl chain length:double bond number) of PS (C) and PE (D) in WT, *ire1*Δ, Δtether, Δtether *ire1*Δ and Δtether *ire1*^W426A^ cells. Data represent mean +/-standard error (N=4 independent experiments). ****, p > 0.0001; ***, p > 0.0002; **, p > 0.0021; *, p < 0.0332; ns, not significant. Abbreviations: CDP, cytidine diphosphate; DAG, diacylglycerol; PA, phosphatidic acid; PC, phosphatidylcholine; PE, phosphatidylethanolamine; PI, phosphatidylinositol; PS, phosphatidylserine; TAG, triacylglycerol

### Ire1 MSR is needed to maintain mono-unsaturated PS and PE in the Δtether cells

As mentioned in the preceding section, the Δtether cells displayed reduced levels of PS and PE (Figure 2B), as observed in previous studies (Quon et al., 2018, Nishimura et al., 2019, Jorgensen et al., 2020, Thomas et al., 2022). Closer examination at the species level (acyl chain length and double bond number) revealed that this was specifically due to significantly decreased levels of mono-unsaturated (32:1 and 34:1) PS and PE in the Δtether cells as compared to wild-type cells (6.2-fold and 3.8-fold, respectively; Figures 2C-D). In contrast, di-unsaturated PS (32:2, 34:2, and 36:2) and PE (primarily 32:2) were not significantly reduced in the Δtether cells (Figures 2C-D). Importantly, the levels of 32:1 and 34:1 PS and PE decreased even further (2.7-fold and 2.5-fold, respectively) in the Δtether *ire1*Δ cells, but not in the Δtether *ire1W426A* cells (Figures 2C-D). Likewise, as compared to Δtether and Δtether *ire1W426A* cells, the Δtether *ire1*Δ cells also displayed further reductions in saturated and mono-unsaturated short acyl chain PE species (26:0, 28:0, 26:1, 28:1, and 30:1) (Figure S2B). Although there were no significant changes in total phosphatidylinositol (PI) levels (Figure 2B) or in the major PI species 32:1 and 34:1 (Figure S2C), the Δtether *ire1*Δ cells displayed statistically significant decreases in saturated and mono-unsaturated short acyl chain PI species (26:0, 28:0, 30:0, and 30:1), as compared to the Δtether and Δtether *ire1W426A* cells (Figure S2C). PC levels were lower in the Δtether cells than wild-type cells, primarily due to decreases in di-unsaturated 32:2 and 34:2 PC, but this effect was independent of Ire1 function (Figures 2B and S2D). Taken together, our lipidomics data indicated that the Ire1 MSR is required to maintain baseline levels of saturated and mono-unsaturated PS, PE, and PI species in the Δtether cells.

### Ire1 MSR modulates sphingolipid metabolism in the Δtether cells

The lipidomics analyses revealed that the Δtether cells have significant drops in PS and PE species containing a saturated C16:0 (palmitate) acyl chain (Figures 2 and S2), which have been reported to be enriched at the PM (Schneiter et al., 1999). We reasoned that the Ire1 MSR might counter this imbalance in GPL homeostasis by regulating other key PM lipids, such as sphingolipids. Indeed, previous studies have reported that Ire1-dependent responses upregulate the expression of sphingolipid synthesis enzymes in yeast and mammalian cells (Travers et al., 2000, Epstein et al., 2012). Likewise, a previous study reported that sphingolipid metabolism is altered in the Δtether cells, including increases in saturated long chain sphingoid bases (Omnus et al., 2016).

Sphingolipid biosynthesis is initiated in the ER via condensation of L-serine and palmitoyl-CoA by the serine palmitoyltransferase complex, termed SPT. The rate-limiting SPT reaction generates 3-ketosphinganine (3-KS) which is then rapidly converted to the long chain base dihydrosphingosine (DHS) (Figure 3A) (Megyeri et al., 2016). DHS is then hydroxylated in yeast by the Sur2 enzyme to form phytosphingosine (PHS) (Haak et al., 1997). To monitor sphingolipid metabolism in the ER of wild-type and mutant cells, we performed pulse labelling experiments using ‘heavy’ ^13^C_3_^15^N_1_-serine followed by targeted mass spectrometry-based lipidomics (Esch et al., 2020, Esch et al., 2023). These experiments revealed significant increases in labelling of the long chain base (LCB) species 18:0 DHS (7-fold), 18:0 PHS (more than 4-fold), and 20:0 PHS (greater than 2-fold) in the Δtether cells as compared to wild-type cells (t = 5 min labelling; Figures 3B-D), consistent with a previous study (Omnus et al., 2016). In contrast, LCB synthesis resembled wild-type rates in the Δtether *ire1*Δ cells (Figures 3B-D). Expression of Ire1^W426A^ in the Δtether *ire1*Δ cells increased LCB labelling, comparable to rates in the Δtether cells (Figures 3B-D). Thus, the Ire1 MSR was sufficient to increase LCB synthesis in the Δtether cells. Accordingly, the Δtether cells displayed resistance to the SPT inhibitor myriocin, but the Δtether *ire1*Δ cells did not (Figure S3), consistent with Ire1-dependent elevations in SPT activity in the Δtether cells.

**Figure 3.**
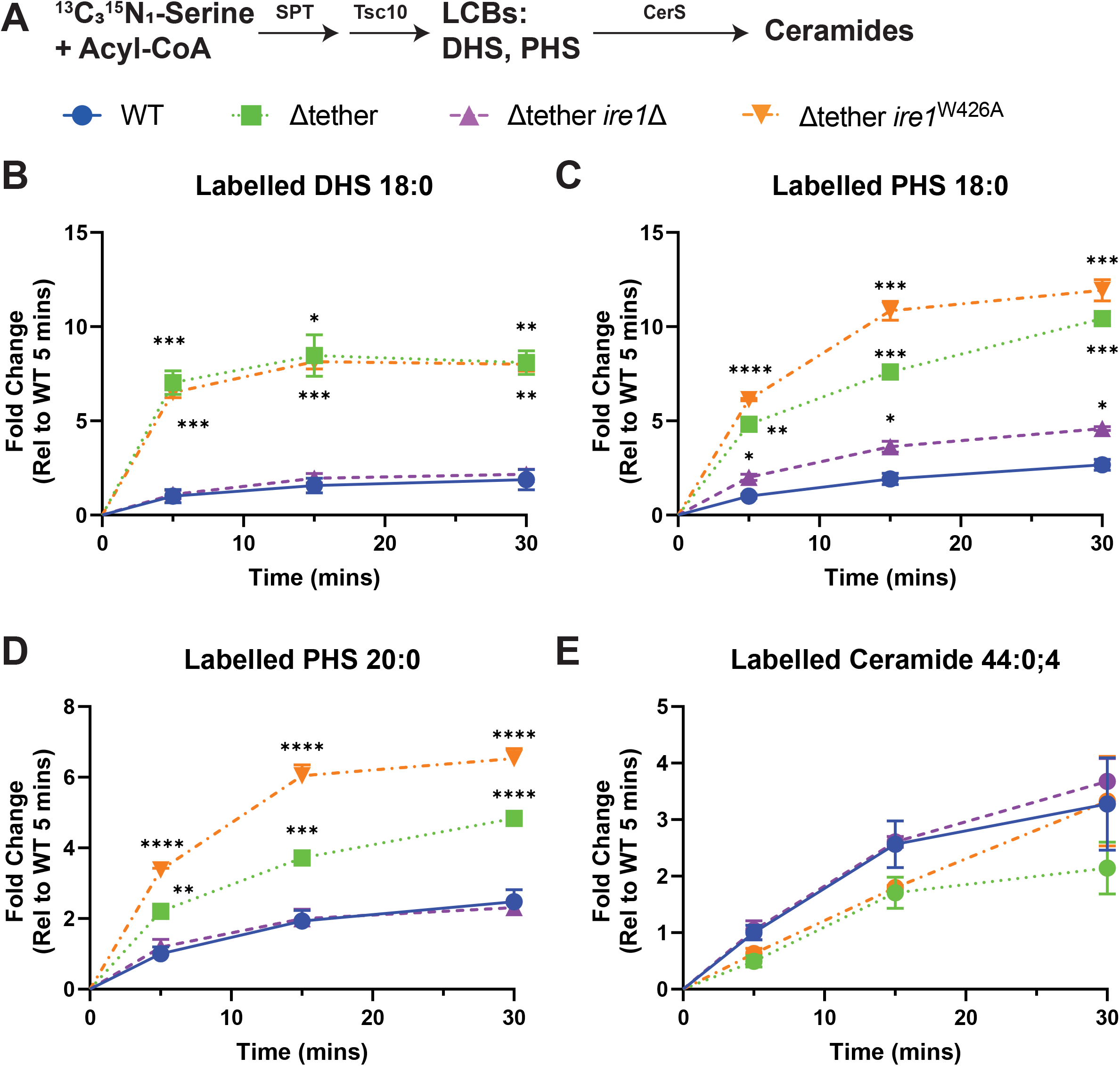
Ire1 MSR regulates sphingolipid synthesis upon loss of ER-PM contacts. (A) Schematic displaying incorporation of ‘heavy’ serine (^13^C^15^N-serine) into synthesis of sphingolipids in the ER. (B, C, D and E) Fold change increases in ‘heavy’ serine-labelled dihydrosphingosine (DHS) 18:0, phytoceramide (PHS) 18:0 and 20:0 and ceramide 44:0 in wild-type (WT), Δtether, Δtether *ire1*Δ, and Δtether *ire1*^W426A^ cells measured after 5, 15, and 30 minutes of labelling with ‘heavy’ serine. Data represent mean +/-standard error (N=3 independent experiments). ****, p > 0.0001; ***, p > 0.0002; ns, not significant.

DHS and PHS are utilized for the generation of dihydroceramide and phytoceramide by the ER-localized ceramide synthases Lac1 and Lag1, respectively (Megyeri et al., 2019) (Figure 3A). Yet, despite increased rates of LCB synthesis in the Δtether cells, the synthesis of ceramides in the ER did not subsequently increase but instead appeared to be slightly impaired in the Δtether and Δtether *ire1W426A* cells, as compared to wild-type cells (1.5-fold to 2-fold at t = 5 and 15 min, although these changes were not statistically significant as analyzed by one-way ANOVA; Figure 3E). Consistent with these results, a previous study reported impaired ceramide synthesis in the Δtether cells (Omnus et al., 2016). Ceramide synthesis rates were similar in wild-type and Δtether *ire1*Δ cells (Figure 3E). Thus, Ire1 MSR promoted LCB synthesis (Figures 3B-D) but not ceramide synthesis (Figure 3E) in the Δtether cells.

We next addressed how SPT-mediated LCB synthesis was increased in the Δtether cells (Figures 3B-D). Previous studies have revealed that the ER membrane-localized Orm1/2 proteins inhibit SPT activity in a ceramide-dependent fashion (Schäfer et al., 2023, Körner et al., 2024, Xie et al., 2024) (see below for further discussion). Thus, reductions in ceramide levels in the ER membrane may bolster SPT-dependent LCB synthesis. Possibly, Ire1 could upregulate the transport of ceramides from the ER to Golgi compartments where they are then converted to the complex sphingolipids inositolphosphorylceramides, IPCs, and mannosylinositol phosphorylceramides, MIPC and M(IP)_2_C (Figure 4A), thereby relieving ceramide-and Orm1/2-mediated inhibition of SPT activity in the ER. To investigate this possibility, we performed ^3^H-myo-inositol labelling experiments and monitored ^3^H-labelled lipids by thin layer chromatography (Figures 4B and S4). If the Ire1 MSR upregulates the transport of ceramides from the ER to Golgi compartments, then synthesis of IPC, MIPC, and M(IP)_2_C may be increased in the Δtether cells, as in the case upon overexpression of the ceramide transfer protein Nvj2 (Liu et al., 2017). However, ^3^H-labelled IPC and M(IP)_2_C were not increased, and MIPC was significantly decreased, in the Δtether cells as compared to wild-type cells (Figure 4B), consistent with a previous study (Omnus et al., 2016). Moreover, if the Ire1 MSR upregulates the conversion of ceramides and PI to IPC, then PI consumption should increase and result in PI depletion (Figure 4A). However, ^3^H-labelled PI and lyso-PI were not decreased in the Δtether cells, indicating that PI consumption needed for IPC synthesis did not increase in the Δtether cells (Figure S4). Consistent with these results, there were no significant differences in the steady-state levels of PI species between the Δtether cells and wild-type cells (Figures 2B and S2B).

**Figure 4.**
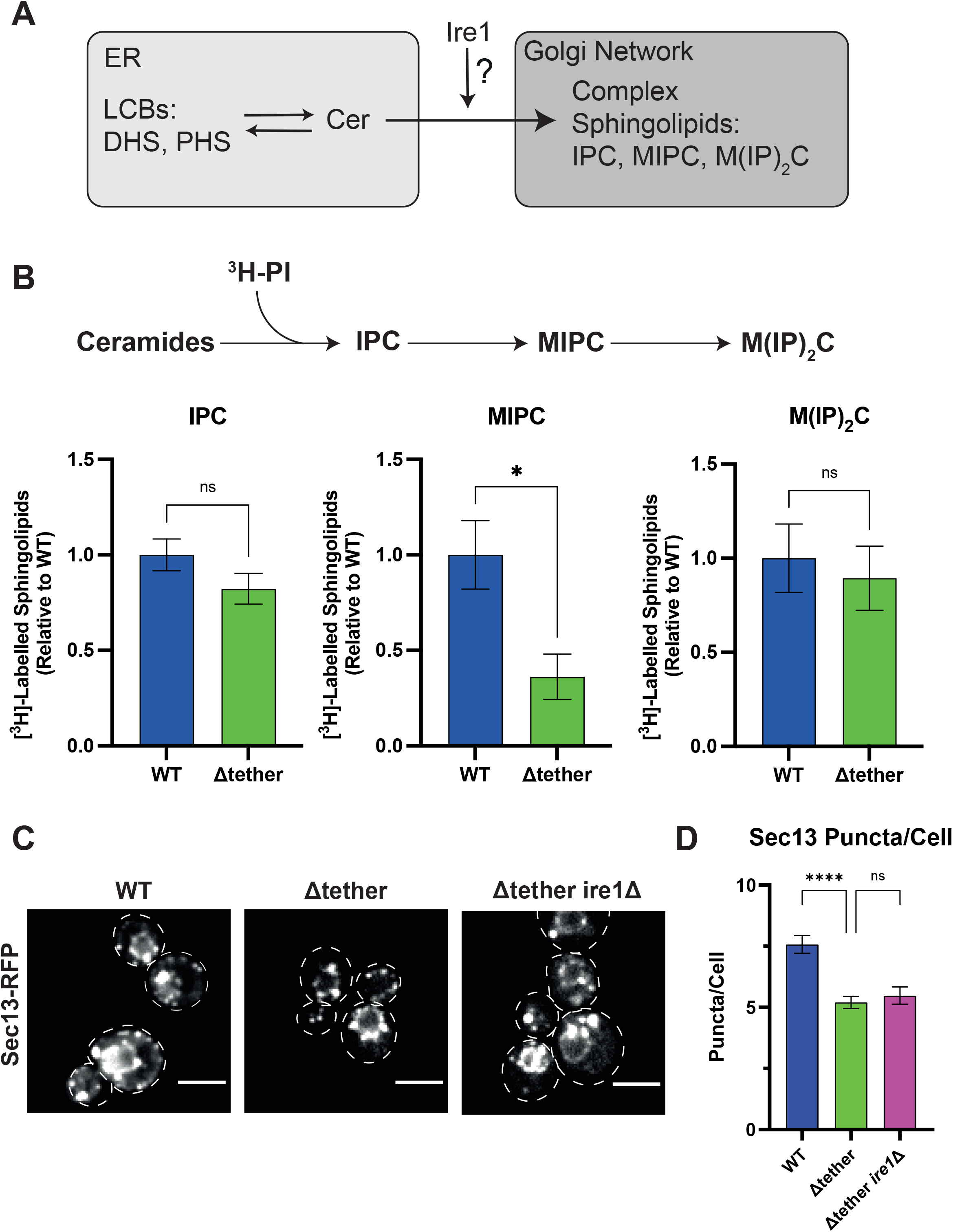
Complex sphingolipid generation and COPII vesicle formation are not increased in the Δtether cells. (A) Schematic overview of sphingolipid metabolism in the ER and Golgi network of budding yeast. The experiments address whether Ire1 promotes export of ceramides from the ER to Golgi compartments and subsequent conversion to complex sphingolipids in the Δtether cells. (B) Top panel: Schematic displaying incorporation of tritiated phosphatidylinositol (^3^H-PI) into complex sphingolipids in the Golgi network. Bottom panel: Measurements of ^3^H-labelled complex sphingolipids in wild-type (WT) and Δtether cells. Data represent mean ± standard error (N=4). * p< 0.033; ns, not significant. C) Representative confocal images of Sec13-RFP in WT, Δtether, and Δtether *ire1*Δ cells. Dashed lines outline cell surfaces. (D) Quantitation of Sec13-RFP puncta in WT, Δtether, and Δtether *ire1*Δ cells. Scale bars, 4μm. Data represent mean +/-standard error (N=3). Raw data are provided in the Data Table. ****, p > 0.0001. Abbreviations: IPC, inositolphosphoceramides; MIPC and M(IP)_2_C, mannosylinositolphosphoceramides

Ceramide lipids traffic from the ER to Golgi compartments via non-vesicular and vesicular transport pathways in yeast, and vesicle-mediated trafficking from the ER is reported to be upregulated upon loss of lipid transport proteins (Kajiwara et al., 2014). However, Sec13-mRFP puncta, corresponding to ER exit sites and COPII vesicles, were decreased in the Δtether cells as compared to wild-type, and loss of Ire1 did not lead to a further reduction in Sec13-mRFP puncta (Figures 4C, D). Intriguingly, immature ER forms of the Gas1 protein were slightly decreased in the Δtether cells as compared to wild-type cells (Figure S4D), suggesting that Ire1 may regulate Gas1 quality control in the ER or trafficking to Golgi compartments in the Δtether cells. Accordingly, loss of Ire1 or Hac1 in the Δtether cells resulted in the accumulation of immature ER Gas1 forms as compared to wild-type cells (Figure S4D). However, expression of Ire1^MFYΔ^ or Ire1^W426A^ did not lower steady-state levels of immature ER Gas1 forms (Figure S4E), while wild-type Ire1 did (Figure S4E), indicating that the UPR was responsible for control of immature ER Gas1 forms in the Δtether cells, consistent with the well-characterized roles of the UPR (Cox and Walter, 1996, Travers et al., 2000, Gardner and Walter, 2011, Walter and Ron, 2011). As such, multiple lines of evidence suggested that the Ire1 MSR did not upregulate ceramide trafficking from the ER in the Δtether cells, and so the Ire1 MSR may regulate sphingolipid metabolism in the ER by some other mechanisms.

### Ire1 regulates Ca^2+^ homeostasis to modulate sphingolipid metabolism in the ER

Previous studies have demonstrated that the yeast ceramide synthases, Lac1 and Lag1, are activated by Ypk1/2-mediated phosphorylation and attenuated by Ca^2+^-and calcineurin-regulated dephosphorylation (Aronova et al., 2008, Muir et al., 2014) (Figure 5A). Likewise, another previous study reported that ceramide synthesis is impaired in the Δtether cells due to increased Ca^2+^-dependent calcineurin activity and hypo-phosphorylation of Lag1 (Omnus et al., 2016). We therefore addressed whether Ire1 may regulate cytosolic Ca^2+^ and calcineurin to modulate ceramide synthesis in the Δtether cells. To examine relative cytoplasmic Ca^2+^ levels in the wild-type and mutant strains, we monitored cytoplasmic GCaMP3 fluorescence by quantitative flow cytometry and confocal microscopy. Cytoplasmic GCaMP3 intensity was approximately 1.8-fold higher in the Δtether cells compared to wild-type (Figure 5B), consistent with previous studies (Omnus et al., 2016, Thomas et al., 2022). In contrast, cytoplasmic GCaMP3 signals were indistinguishable from wild-type in the Δtether *ire1*Δ cells (Figure 5B). We also examined relative Ca^2+^ levels in the ER lumen and found that the intensity of a GCaMP3-HDEL reporter was nearly 3-fold higher in the Δtether cells as compared to wild-type (Figure S5). GCaMP3-HDEL signals were significantly lower in the Δtether *ire1*Δ cells as compared to the Δtether cells (Figure S5). Altogether, these results suggested that Ire1-dependent responses resulted in elevated Ca^2+^ levels in the cytoplasm and ER upon loss of ER-organelle contacts.

**Figure 5.**
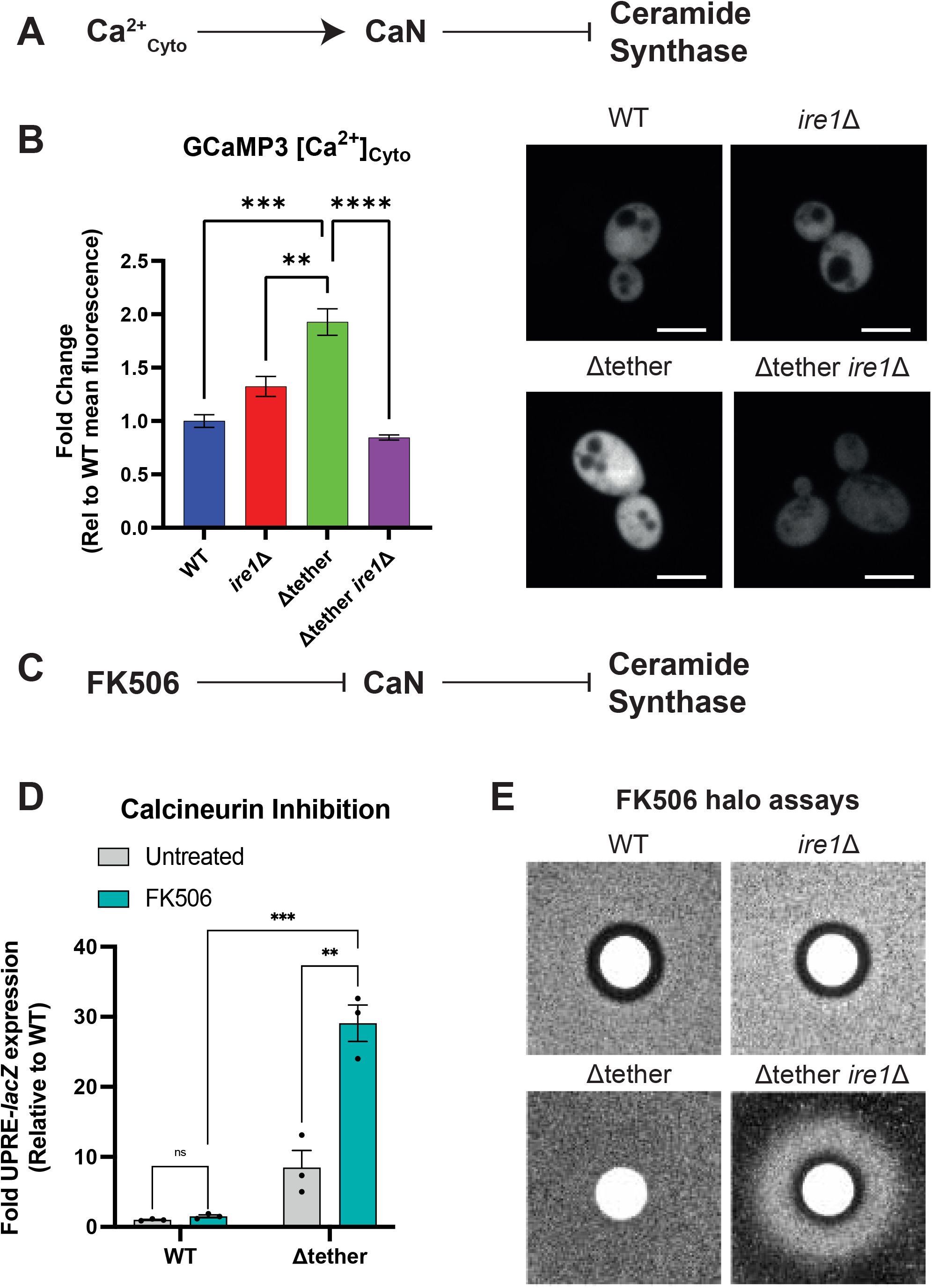
Ire1 regulates Ca^2+^ and calcineurin to maintain ER homeostasis. (A) Ca^2+^ and calcineurin (CaN) attenuate ceramide synthases in budding yeast (Aronova et al., 2008, Muir et al., 2014, Omnus et al., 2016). (B) Left panel: Mean fluorescence intensity of cytoplasmic GCaMP3 Ca^2+^ reporter in wild-type (WT), Δtether, Δtether *ire1*Δ, and *ire1*Δ cells as measured by flow cytometry. Data represent mean ± standard error (N=5 experiments, n=100,000 cells per experiment). Right: Confocal imaging of the cytoplasmic GCaMP3 Ca^2+^ reporter in WT, Δtether, Δtether *ire1*Δ and *ire1*Δ cells. Scale bar, 4 μm. (C) FK506 inhibits CaN activity that in turn inhibits ceramide synthesis in budding yeast. (D) Inhibition of CaN with FK506 induces robust expression of the *UPRE-lacZ* reporter in the Δtether cells. Data represents mean +/-standard error (N=4). (E) WT, Δtether, and Δtether *ire1*Δ cells were plated as lawns in agar and then overlaid with sterile filter discs containing 10 μl of a 10 mg/ml stock of FK506 in DMSO. After incubation at 30°C for 3 days, the plates were imaged. WT and *ire1*Δ cells displayed clear zones of growth inhibition (halos), whereas the Δtether cells did not, consistent with increased CaN activity in the Δtether cells, as previously reported (Omnus et al., 2016). The Δtether *ire1*Δ cells showed an initial zone of growth inhibition (halo) like WT cells, indicating that Ire1 is essential for FK506 resistance in the Δtether cells. Intriguingly, cell growth improved and then declined as FK506 concentration decreased, suggesting that the Δtether *ire1*Δ cells require a window of optimal CaN activity for growth. ****, p < 0.0001; ***, p < 0.0002; *, p < 0.0332.

Next, we addressed the effects of impairing Ca^2+^-dependent calcineurin on ER stress responses in the Δtether cells. Treatment of wild-type cells with the calcineurin inhibitor FK506 (Figure 5C) (Liu et al., 1991) did not induce significant expression of the *UPRE-lacZ* reporter (Figure 5D). However, inhibition of calcineurin with FK506 resulted in a striking increase in *UPRE-lacZ* expression in the Δtether cells (nearly 30-fold above wild-type basal levels, Figure 5D), resembling tunicamycin-induced Ire1 UPR (see Figure 1C). Because a previous study reported that calcineurin activity is increased in the Δtether cells (Omnus et al., 2016), we next addressed whether loss of Ire1 impacts calcineurin activity in the Δtether cells. While wild-type cells displayed sensitivity to FK506 as assessed by zones of growth inhibition in halo assays, growth of the Δtether cells was not impeded by FK506 (Figure 5E), consistent with previous findings (Omnus et al., 2016). In contrast, the Δtether *ire1*Δ cells were sensitive to FK506 and formed initial zones of growth inhibition (Figure 5E) The Δtether *ire1*Δ cells also displayed a more complex growth pattern in the halo assays, where an optimal range of FK506-sensitive calcineurin activity permitted robust growth (Figure 5E). Altogether, these results suggest that Ire1-dependent Ca^2+^ signals upregulate calcineurin activity in the Δtether cells (Figures 5B and 5E) and that inhibition of calcineurin in the Δtether cells resulted in severe ER homeostasis defects (Figure 5D).

### Ire1 controls steady-state levels of DHS and ceramides in the in the Δtether cells

As mentioned, sphingolipid biosynthesis is initiated in the ER membrane via the serine palmitoyltransferase (SPT) complex that consists of the integral ER membrane proteins Lcb1, Lcb2 and Tsc3. The SPT subunits associate with additional integral ER membrane proteins including Orm1 and Orm2 that negatively regulate SPT activity in a ceramide-dependent manner (termed the SPOT complex) (Breslow et al., 2010, Schäfer et al., 2023, Körner et al., 2024, Xie et al., 2024), providing a negative feedback loop to prevent the build-up of LCBs and ceramides. Our results revealed that the Ire1 MSR increases SPT-dependent LCB synthesis but not ceramide synthesis (Figure 3), suggesting that Ire1 MSR prevents the accumulation of ceramides. To address this possibility, we determined the relative steady-state levels of ceramides, as well as LCBs, in wild-type, Δtether, Δtether *ire1*Δ, and Δtether *ire1W426A* cells by mass spectrometry-based lipidomics. As expected, ceramide levels were significantly lower (2-fold) in the Δtether and Δtether *ire1W426A* cells, but not Δtether *ire1*Δ cells, as compared to wild-type cells (Figure 6B). Unexpectedly, even while LCB synthesis rates were elevated in the Δtether cells (Figure 3), there were no significant changes in the steady-state levels of PHS (the major LCB in yeast) in any of the strains tested (Figure 6C). Moreover, DHS levels were reduced in the Δtether cells (<60% of wild-type levels) and decreased even further in the Δtether *ire1*Δ cells (<25% of wild-type levels) (Figure 6D). DHS levels were not significantly different in the Δtether cells and Δtether *ire1W426A* cells (Figure 6D). Thus, Ire1 was required for increased DHS synthesis as well as sustaining DHS steady-state pools in the Δtether cells (Figures 3B and 6D). We also addressed whether Ire1 responses might regulate levels of the Orm1/2 proteins in the Δtether cells. As monitored by quantitative confocal microscopy, GFP-Orm1 and GFP-Orm2 fluorescence intensities were significantly higher in the Δtether *ire1*Δ cells (>3-fold and >5-fold, respectively, as compared to wild-type cells, Figure S6). Thus, in the Δtether cells, Ire1 responses are important to restrict steady-state levels of ceramides and the Orm1/2 proteins, as well as to maintain dynamic pools of DHS.

**Figure 6.**
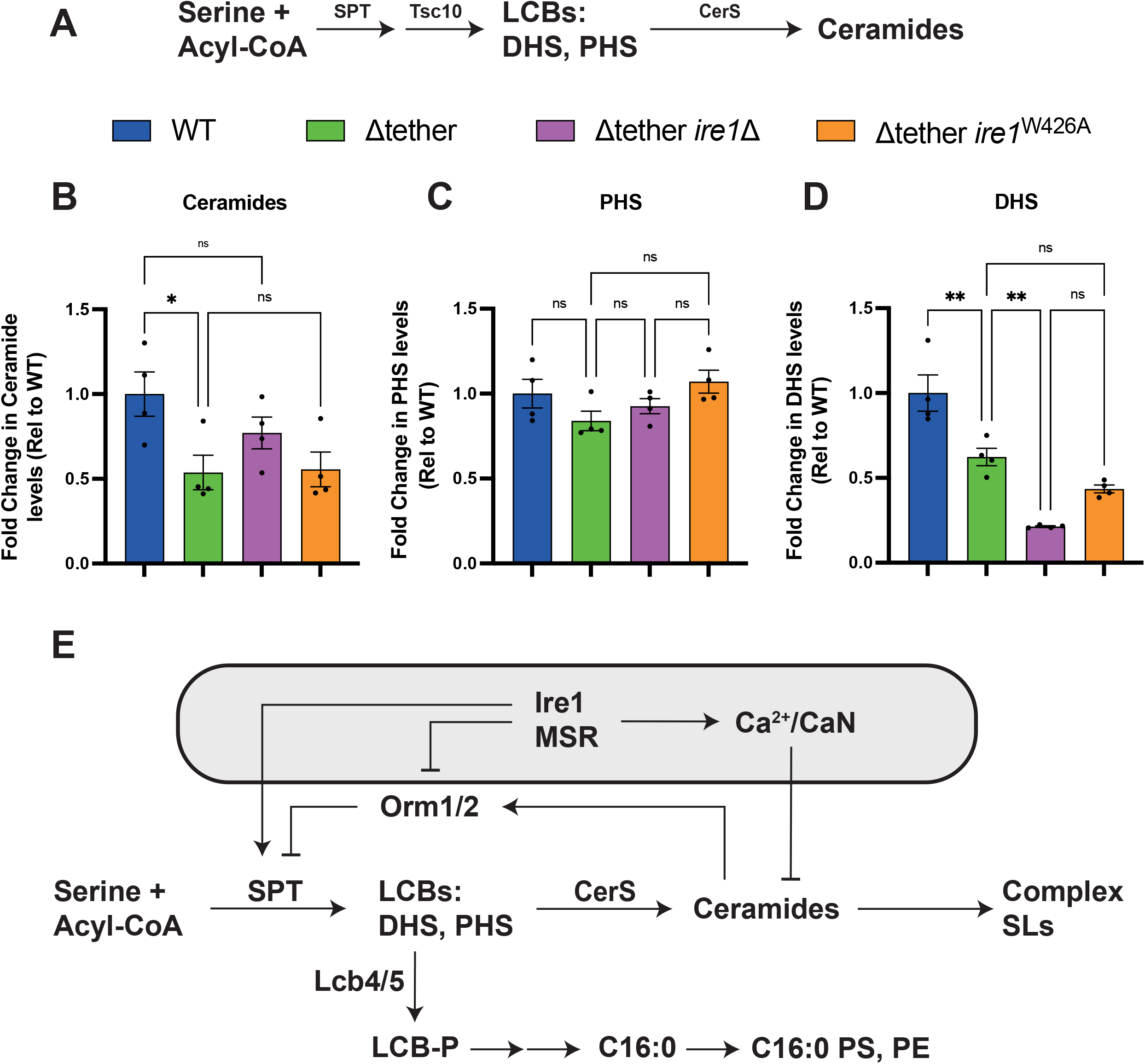
Ire1 controls steady-state levels of DHS and ceramides in the Δtether cells. (A) Schematic representation of sphingolipid synthesis in the ER of budding yeast. (B-D) Relative fold changes in steady-state levels of ceramides (B), PHS (C), and DHS (D) in wild-type (WT), Δtether, Δtether *ire1*Δ, and Δtether *ire1*^W426A^ cells. Data represent mean +/-standard error (N=4). ****, p > 0.0001; ***, p > 0.0002; ns, not significant. (E) Potential roles of the Ire1 MSR upon disruptions in saturated GPL synthesis in the Δtether cells. Ire1 MSR upregulates Ca^2+^-dependent CaN activity (1), which keeps ceramide levels in the ER in check (2), thereby preventing ceramide-and Orm1/2-mediated inhibition of SPT activity as well as ceramide-induced cell death (3). Ire1 also upregulates expression of the SPT subunit Lcb1 (Travers et al., 2000). LCB synthesis may support cell growth directly or via increased production of LCB phosphates, complex sphingolipids, or a shunt pathway that generates saturated fatty acids for GPL synthesis. See Discussion for further details. Abbreviations: C16:0, palmitate; CerS, ceramide synthase; DHS, dihydrosphingosine; GPL, glycerophospholipid; LCBs, long chain bases; LCB-P, LCB phosphates; MSR, membrane stress response; PHS, phytosphingosine; PE, phosphatidylethanolamine; PS, phosphatidylserine; SLs, sphingolipids; SPT, serine palmitoyl transferase

Collectively, our results indicate that impaired lipid transport out of the ER results in reductions in mono-unsaturated (C16:0 acyl chain-containing) PS and PE species in the Δtether cells (Figure 2 and S2), and that the Ire1 MSR upregulates the production of saturated LCBs (DHS and PHS) (Figure 3), possibly to maintain lipid acyl chain saturation homeostasis which is critical in the control of the biophysical properties of membrane bilayers. The data further suggest that the Ire1 MSR prevents ceramide-and Orm1/2-mediated inhibition of SPT (Figures 6 and S6). Intriguingly, while DHS and PHS synthesis are increased by the Ire1 MSR (Figure 3), steady-state levels of PHS and DHS are not elevated, suggesting these LCBs (especially DHS) may be converted to other lipids, possibly mono-unsaturated PS and PE species, rather than ceramides (Figures 2, 3, and 6B), that are essential for normal cell growth (Figure 6E and see Discussion).

## Discussion

Numerous studies have established that membrane contact sites direct lipid metabolism and non-vesicular transport of lipids from the ER (Nishimura and Stefan, 2020, Prinz et al., 2020, Voeltz et al., 2024). In this study, we have addressed how disruptions in membrane contact sites impact ER stress responses and their roles in membrane lipid homeostasis. Using Δtether yeast cells as a model system, we found that Ire1-mediated membrane stress responses (MSR) were necessary and sufficient for cell growth upon loss of membrane contact sites. Importantly, our findings further revealed that the Ire1 MSR confers essential roles in sustaining threshold glycerophospholipid levels as well as modulating sphingolipid synthesis in the Δtether cells. These findings contrast with previous studies which concluded that Ire1 does not adjust lipid metabolism upon disruptions in glycerophospholipid synthesis (Thibault et al., 2012, Ho et al., 2020).

### Ire1 senses membrane bilayer stress in the Δtether cells

A previous study reported constitutive Ire1 activity in the Δtether cells (Manford et al., 2012). However, the nature of ER stress experienced in the Δtether cells was not investigated. Our lipidomic analyses confirmed decreases in steady-state levels of the glycerophospholipids PS, PE, and PC in the Δtether cells (Figures 2 and S2). Accordingly, alterations in glycerophospholipid metabolism have been reported to result in membrane bilayer stress and Ire1 activation (Jonikas et al., 2009, Promlek et al., 2011, Thibault et al., 2012, Halbleib et al., 2017, Ho et al., 2020). In particular, the Ire1 transmembrane and juxtamembrane regions have been proposed to induce compression of the ER bilayer (Halbleib et al., 2017). Consequently, increases in ER bilayer thickness and packing order (governed by acyl chain length and saturation), which oppose ER bilayer compression, have been shown to promote the association of Ire1 dimers and Ire1 activation, presumably within remnant ER membrane subdomains that are thinner, more disordered, and less resistant to bilayer compression (Pineau et al., 2009, Han et al., 2010, Promlek et al., 2011, Surma et al., 2013, Volmer and Ron, 2015, Halbleib et al., 2017, Rajakumar et al., 2024). The glycerophospholipid saturation index did not increase in the Δtether cells, and instead there were significant reductions in species containing a saturated C16:0 acyl chain (Figures 2 and S2). Possibly, a subtle increase in ER bilayer thickness (due to decreases in C32 and C34 species but not C36 species) may account for Ire1 activation in the Δtether cells. In addition, impaired transport of LCBs and ceramides with long, saturated acyl chains out the ER may trigger Ire1 activation in the Δtether cells. Previous studies have implicated the tricalbin proteins (Tcb1/2/3 which are missing in the Δtether cells) in the transport of these sphingolipids from the ER (Ikeda et al., 2020, Hanaoka et al., 2024). The Δtether cells also demonstrated mild proteotoxic stress as well as lipid bilayer stress (Figures 1 and S1). Because perturbations in membrane lipid homeostasis can induce potentially toxic protein misfolding and degradation (Shyu et al., 2019, Renne and Ernst, 2023, Ernst et al., 2024), the primary role of the Ire1 MSR may be to alleviate membrane bilayer stress by modulating membrane lipid metabolism, in order to prevent severe proteostasis defects and chronic ER stress.

### Ire1 MSR confers essential roles in lipid metabolism

Ire1 and Hac1 have been shown to upregulate the expression of genes that encode components of COPII vesicles, including Sfb2 and Sfb3 (Travers et al., 2000). Thus, we addressed whether Ire1 upregulated vesicle-mediated trafficking of lipids from the ER. However, Sec13 puncta were significantly decreased in the Δtether cells (Figures 4C and 4D), arguing against an essential role for Ire1 in upregulation of COPII vesicle-mediated lipid trafficking from the ER in the Δtether cells. Ire1 and Hac1 have also been reported to upregulate the expression of genes that encode glycerophospholipid biosynthesis enzymes (Travers et al., 2000), including Opi3 which is involved in PC synthesis (Greenberg et al., 1983). However, there was no significant difference in steady-state PC levels between Δtether and Δtether *ire1*Δ cells (Figure 2B), ruling out an essential role for Ire1 in PC synthesis, consistent with previous work demonstrating that PC synthesis is not essential for growth of yeast cells (Bao et al., 2021). Instead, lipidomics data revealed that the Ire1 MSR was required to maintain threshold levels of mono-unsaturated PS and PE species (Figures 2C and 2D), as well as PI species with saturated short acyl chains (Figure S2C). Thus, the requirement for Ire1 MSR in the Δtether cells correlated with homeostatic regulation of saturated and mono-unsaturated glycerophospholipids.

Mono-unsaturated PS and PE species have been shown to be enriched at the PM in yeast cells (Schneiter et al., 1999). As a compensatory mechanism for decreased mono-unsaturated PS and PE, we addressed whether the Ire1 MSR upregulated the synthesis of other saturated lipids, such as sphingolipids, to support cell growth. Consistent with this notion, previous studies have indicated that Ire1 upregulates sphingolipid synthesis in yeast (Travers et al., 2000, Epstein et al., 2012). Accordingly, ^13^C_3_^15^N_1_-serine metabolic labelling experiments coupled with targeted lipidomics revealed increased rates of LCB synthesis in the Δtether cells (Figures 3B-D). This result was also surprising, as a previous study found that SPT activity is highest in the cortical ER network (Esch et al., 2023), which is depleted in the Δtether cells (Manford et al., 2012). However, the Ire1 MSR (conferred by Ire1^W426A^) was necessary and sufficient to drive LCB synthesis upon loss of the cortical ER network (Figures 3B-D). Taken together, our findings suggested that Ire1 MSR redirects the metabolic channeling of serine and palmitoyl-CoA toward the synthesis of saturated LCBs (Figure 3) upon decreases in mono-saturated PS species the Δtether cells (Figure 2).

LCBs are converted to dihydroceramides and phytoceramides in the ER by the ceramide synthases Lac1 and Lag1 in budding yeast (Figure 3A). Remarkably, while LCB synthesis was significantly increased in the Δtether cells, ceramide synthesis did not subsequently increase (Figure 3E). Likewise, steady-state levels of ceramides were significantly decreased in Δtether and Δtether *ire1*^W426A^ cells as compared to wild-type cells, but not in the Δtether *ire1*Δ cells (Figure 6). Excessive ceramides can result in membrane leaflet condensations (extremes in leaflet packing due to the ability of ceramides to undergo extensive hydrogen bonding and their saturated acyl chains) and subsequent perturbations in lateral bilayer organization (Lopez-Montero et al., 2010). Imbalances in ceramides across membrane leaflets can also induce normally rare lipid ‘flip-flop’ events that can lead to bilayer permeability defects (Contreras et al., 2009). Consequently, ceramides must be precisely controlled and elevations in ceramides often result in lipotoxicity and loss of cell viability (Barisch et al., 2023). Multiple lines of evidence have shown that Ire1 is highly tuned to respond to increases in LCBs and ceramides in the ER (Pineau et al., 2009, Han et al., 2010, Rajakumar et al., 2024) (Figure 1).

### Ire1 MSR controls sphingolipid metabolism by regulating Ca^2+^, calcineurin, and Orm1/2 levels

Our results further revealed that Ire1 MSR promoted LCB synthesis while preventing ceramide accumulation (Figures 3 and 6). Nvj2 has been implicated as an ER stress-induced ceramide transfer protein, but Nvj2 activity was reportedly independent of Ire1 (Liu et al., 2017). Accordingly, our data did not support increased transport of ceramides from the ER to Golgi compartments in the Δtether cells (Figure 4). Instead, the conversion of LCBs to ceramides may be held in check in the Δtether cells, potentially due to increased cytoplasmic Ca^2+^ and calcineurin phosphatase activity (Figure 5) (Omnus et al., 2016), which has been shown to attenuate the yeast ceramide synthases Lac1 and Lag1 (Aronova et al., 2008, Muir et al., 2014). ER stress has been reported to activate the high-affinity Ca^2+^ influx system (HACS), albeit in an Ire1-independent manner (Bonilla et al., 2002). However, a subsequent study found that HACS activation depended on potassium influx via Kch1, Trk1 and Trk2 (Stefan and Cunningham, 2013). Importantly, both Kch1 and Trk2 have been reported to be upregulated by Ire1 (Travers et al., 2000, Stefan and Cunningham, 2013), suggesting Ire1 may contribute to elevated cytoplasmic Ca^2+^ upon membrane bilayer stress.

Ire1-mediated HACS activity may trigger the tricalbin proteins (Tcb1/2/3), shown to function upon elevations in cytoplasmic Ca^2+^ (Thomas et al., 2022), to transport ceramides from the ER (Ikeda et al., 2020, Hanaoka et al., 2024). However, the Δtether cells lack Tcb1/2/3 and therefore must rely on additional homeostatic mechanisms to resolve ER membrane stress. Importantly, inhibition of the Ca^2+^-activated calcineurin phosphatase resulted in severe ER stress (Figure 5), consistent with previous work indicating a role of calcineurin in resolving ER stress (Bonilla et al., 2002). Interestingly, calcineurin also regulates the expression of the Ca^2+^ ATPases Pmc1 and Pmr1, which transport Ca^2+^ from the cytoplasm into the vacuole and secretory compartments (Matheos et al., 1997, Stathopoulos and Cyert, 1997), and are necessary to prevent cytotoxic Ca^2+^ overload (Cunningham and Fink, 1994). Pmr1 may account for increased Ca^2+^ signals in the ER of the Δtether cells (Figure S5). Thus, in response to ER membrane stress in yeast, an Ire1-dependent regulatory circuit triggers Ca^2+^ influx via HACS, subsequent Ca^2+^-and calcineurin-mediated inhibition of ceramide synthesis to prevent ceramide lipotoxicity, as well as calcineurin-regulated transport of Ca^2+^ into cellular stores to prevent cytotoxic Ca^2+^ overload.

Similar homeostatic responses exist in mammalian cells and result in disease when disrupted. For instance, IRE1α has been implicated in store-operated Ca^2+^ entry (SOCE) (Carreras-Sureda et al., 2023). While mammalian ceramide synthases may not be subject to calcineurin-mediated inhibition, calcineurin has been shown to inhibit neutral sphingomyelinase that generates ceramide as its product (Filosto et al., 2010), which may be important in preventing ceramide lipotoxicity. In turn, IRE1α and XBP1 have been found to control the expression of the sarco/endoplasmic reticulum Ca^2+^ ATPase isoform SERCA2b that transports Ca^2+^ from the cytoplasm into the ER (Park et al., 2010), and SERCA2b dysfunction has been implicated in motor neuron disease (Saito et al., 2022). Accordingly, SERCA isoforms require specific membrane lipid environments to function (Gustavsson et al., 2011), and dysregulation of ER membrane lipid homeostasis results in loss of SERCA function (Li et al., 2004, Fu et al., 2011).

Consequently, impairment of IRE1α and XBP1 function that occurs in familial ALS8 due to the VAP-B P56S mutant protein (Kanekura et al., 2006, Suzuki et al., 2009, Chen et al., 2010), along with ER membrane lipid and Ca^2+^ homeostasis defects, may result in a vicious cycle that contributes to motor neuron degeneration. Interventions that safely install the adaptive MSR, restoring Ca^2+^ and membrane lipid homeostasis, may be protective and prevent disease progression.

Our results also revealed roles of Ire1 in the regulation of steady-state levels of ceramides and the Orm1/2 proteins which bind ceramides to inhibit SPT activity in the ER (Schäfer et al., 2023, Xie et al., 2024). Increased levels of the Orm1/2 proteins, along with elevations in ceramides (Figures 6 and S6), may explain why SPT-dependent LCB synthesis was impaired in the Δtether *ire1*Δ cells as compared to the Δtether cells (Figure 3). Intriguingly, Orm2 steady-state levels have been shown to increase upon ER protein misfolding and trafficking defects (Gururaj et al., 2013, Platzek et al., 2025). However, ER stress-induced Orm2 expression was reported to be calcineurin-dependent and Ire1-independent (Gururaj et al., 2013). Even if calcineurin activity boosts Orm2 expression, this may not result in SPT inhibition because calcineurin has been shown to effectively attenuate ceramide synthesis (Aronova et al., 2008, Muir et al., 2014, Omnus et al., 2016). Orm2 protein levels have been shown to be regulated primarily by the endosome-and Golgi-associated degradation (EGAD) pathway (Schmidt et al., 2019). Conceivably, impaired ER to Golgi trafficking of Orm2 may impair EGAD-mediated downregulation of the Orm2 protein. Calcineurin has been shown to antagonize EGAD-mediated Orm2 downregulation (Schmidt et al., 2019). Under ER stress conditions, Orm2 may then become subject to the ER-associated degradation (ERAD) pathway which has been shown to be regulated by Ire1 (Travers et al., 2000). Accordingly, control of Orm2 levels in the ER may become highly dependent on Ire1-regulated ERAD in the Δtether cells which display reduced COPII vesicle formation (Figure 4) and increased calcineurin activity (Figure 5) (Omnus et al., 2016). Likewise, Ire1 was essential for quality control of the GPI-anchor protein Gas1 in the Δtether cells (Figure S4). Thus, Ire1 responses are immensely nuanced in their sensitivity, plasticity, and capacitive functions. To acutely resolve membrane stress, Ire1-mediated Ca^2+^ signals and calcineurin activity attenuate ceramide synthesis. Upon prolonged stress conditions where persistent calcineurin activity may become problematic, potentially due to increasing Orm2 (Gururaj et al., 2013, Schmidt et al., 2019), Ire1 prevents toxic Orm2 accumulation to avoid shut down of sphingolipid synthesis in the ER.

Our findings indicated that the Ire1 MSR upregulated LCB synthesis upon disruptions in glycerophospholipid metabolism (Figures 2 and 3). However, steady-state LCB levels were not increased in the Δtether cells (Figure 6). Specifically, Ire1 MSR was required for increased DHS synthesis (Figure 3B) as well as to maintain dynamic pools of DHS (Figure 6D). Thus, DHS may be converted to lipids that are essential for growth (Figure 6E). Previous studies have elucidated a shunt pathway whereby phosphorylated LCBs (LCB-P) are converted to saturated C15 and C16 fatty acids that can be utilized for glycerophospholipid synthesis (Funato et al., 2003, Nakahara et al., 2012, Kondo et al., 2014). A key step in this pathway involves the LCB kinase Lcb4 that generates LCB-P species (Figure 6E), and loss of Lcb4 has been shown to impair glycerophospholipid synthesis (Funato et al., 2003, Kondo et al., 2014). Thus, the Ire1 MSR may upregulate synthesis of saturated LCBs to sustain baseline levels of mono-unsaturated PS and PE (Figures 2, 3, and 6E). Consistent with this notion, *LCB4* was identified as a candidate hit in a synthetic lethality genetic screen using the Δtether cells (Jorgensen et al., 2020). We also found that exogenous addition of PHS relieved ER stress in the Δtether cells (Figure 1B). Accordingly, Ire1 MSR not only prevents ceramide lipotoxicity but also ensures the synthesis of threshold pools of critical lipid species (Figure 6E). Thus, the Ire1 MSR directs multiple homeostatic regulatory mechanisms which adjust metabolic lipid channeling upon disruptions in glycerophospholipid synthesis, in contrast with previous claims (Thibault et al., 2012, Ho et al., 2020).

### Targeting Adaptive IRE1α Responses to Prevent Motor Neuron Degeneration in ALS8

Our findings from yeast suggest that targeting adaptive ER stress responses may be beneficial in diseases caused by defects in inter-organelle contact proteins. For example, Amyotrophic Lateral Sclerosis type 8 (ALS8) is a rare motor neuron disease caused by a proline-to-serine substitution at position 56 in the VAP-B protein (Nishimura et al., 2005). The ALS8-associated P56S substitution results in misfolding of the VAP-B protein and subsequent alterations in ER morphology and homeostasis (Borgese et al., 2021). Intriguingly, the IRE1α-and ATF6-mediated branches of the UPR are impaired in VAP-B P56S-expressing cells, whereas PERK1-mediated cell death appears to be upregulated (Kanekura et al., 2006, Gkogkas et al., 2008, Suzuki et al., 2009, Chen et al., 2010, Landry et al., 2025). Thus, combined defects in ER homeostasis and failure to mount protective ER stress responses may contribute to motor neuron degeneration during the progression of ALS8. In support of this notion, recent studies have suggested that enforced expression of XBP1s, the spliced form of XBP1 mRNA, may be protective in model ALS systems (Li et al., 2025, Valenzuela et al., 2025). However, extreme caution is warranted as chronic, maladaptive ER stress responses result in cellular degeneration (Walter and Ron, 2011). Indeed, another recent study reported that inhibition of the PERK branch of the UPR, a driver of cell death upon chronic ER stress (Walter and Ron, 2011), rescues phenotypes in inducible pluripotent stem cell-derived neurons expressing VAP-B P56S (Landry et al., 2025). Potentially, combinatorial approaches that target adaptive IRE1α responses and attenuate maladaptive PERK cell death may be synergistically beneficial in treating ALS and possibly other neurodegenerative diseases caused by imbalances in membrane lipid homeostasis.

## Materials and Methods

### Yeast strains, plasmids, and growth assays

Descriptions of yeast strains, plasmids, reagents, and resources used in this study are listed in the Reagents Table. Gene deletions were introduced into yeast by homologous recombination (Longtine et al., 1998). Standard techniques and media were used for yeast and bacterial growth. For plasmid shuffle growth assays (Figures 1E and S1A), cells were grown to mid-log phase, adjusted to 1 OD_600_/ml, and ten-fold serial dilutions were plated on agar media either lacking uracil and leucine (to maintain the *URA3*-marked and *LEU2*-marked *IRE1* plasmids) or lacking leucine and containing 5-fluoroorotic acid (5-FAO, to select against the *URA3*-marked *IRE1* plasmid). To monitor zones of growth inhibition (halo assays, Figures S3 and 5E), lawns of the indicated yeast strains were plated in soft agar and overlaid with sterile filters containing the SPT inhibitor myriocin (3 μl of a 1 mg/ml stock of myriocin in DMSO, Figure S3) or the calcineurin inhibitor FK506 (10 μl of a 10 mg/ml stock of FK506 in DMSO). The plates were imaged after incubation at 30°C for 3 days.

### *UPRE-lacZ* β-galactosidase assays

The indicated strains harbouring a *UPRE-lacZ* reporter (Cox and Walter, 1996) were grown to mid-log phase and treated with 50 μg/ml PHS, 100 μg/ml C8-Ceramide, 2 μg/ml tunicamycin (Figures 1B and 1C), or 100 μg/ml FK506 (Figure 5D). Cells were then harvested by centrifugation, washed in Z buffer (60 mM Na_2_HPO_4_, 40 mM NaH_2_PO_4_), 10 mM KCl, 1 mM MgSO_4_), and the OD_600_ was measured. Samples were then adjusted to 500μl in Z buffer, and 50 μl 0.1% (w/v) SDS was added. Samples were vortexed for 15 seconds, 50 μl chloroform was added, and samples were vortexed again for 15 seconds. 100 μl of the β-galactosidase substrate ONPG (4 mg/ml stock) was added to start the reaction, which was quenched by adding 500 μl 1M Na_2_CO_3_. Samples were spun at 13,000 x *g* for 2 minutes. The absorbance (Abs_420_) of the supernatant was measured, and the units of activity were calculated using the equation: (1000 x Abs_420_)/time (min) x OD_600_). The raw data for the *UPRE-lacZ* assays are provided in the Data Table.

### Cell growth and sample preparation for mass spectrometric measurements of glycerolipids, LCBs, and ceramides

The indicated yeast strains (carrying either an empty *LEU2*-marked plasmid or *LEU2*-marked *ire1W426A* plasmid) were inoculated from a logarithmic growing preculture and then grown to exponential growth phase at 30°C in YND lacking leucine. Cells were collected at 2250 x *g* for 5 min at 4°C and snap frozen in liquid nitrogen. Cells were lysed with glass beads in 500 μl 155 mM ammonium formate using a FastPrep machine (MP Biomedicals). Cell lysates were adjusted to 200 μg of protein (total) and lipids were extracted using 2:1 chloroform/methanol (Ejsing et al., 2009). Prior to extraction, an internal standard mixture containing EquiSPLASH, sphingosine d17:1 and ceramide d17:1/24:0 (Avanti Polar Lipids) was added to each sample for normalization. Dried lipid samples were dissolved in a 65:35 mixture of mobile phase A (50:50 water/acetonitrile, including 10 mM ammonium formate and 0.1% formic acid) and mobile phase B (88:10:2 2-propanol/acetonitrile/H_2_0, including 2 mM ammonium formate and 0.02% formic acid). Samples were subjected to LC-MS/MS analysis to determine glycerophospholipid levels or targeted analysis of LCB and ceramide levels, as described below.

The metabolic flux analysis of LCBs and ceramides was performed as previously described (Esch et al., 2020, Esch et al., 2023). The indicated yeast strains (carrying either an empty *LEU2*-marked plasmid or *LEU2*-marked *ire1W426A* plasmid) were inoculated from a logarithmic growing preculture in 20 ml YND lacking leucine and grown to exponential growth phase at 30°C until they reached an OD_600_ of 0.8. [^13^C_3_^15^N_1_]-serine (CCN3000P1; CortecNet) was added to a final concentration of 3.8 mM. Samples (2.5 OD_600_ units each) were collected after 5, 15 and 30 min (t=5, 15, 30) at 2250 x *g* for 2 min at 4°C and cell pellets were directly snap frozen in liquid nitrogen. Lipids were extracted using 2:1 chloroform/methanol (Ejsing et al., 2009). Prior to extraction, an internal standard mixture containing sphingosine d17:1 and ceramide d17:1/24:0 (Avanti Polar Lipids) was added to each sample for normalization. Dried lipid samples were dissolved in a 65:35 mixture of mobile phase A and mobile phase B. Samples were analysed as described in the section targeted analysis of LCBs and ceramides by LC-MS/MS. All LC-MS/MS lipidomic raw data are provided in the Data Table.

### Measurements of glycerolipid levels by liquid chromatography-tandem mass spectrometry (LC-MS/MS) analysis

The LC-MS/MS analysis of glycerolipids (GPLs, DAG, TAG) was performed as previously described (Esch et al., 2020, Esch et al., 2023). Extracted lipids were subjected to HPLC analysis employing a C18 reverse-phase column (Thermo Accucore RP-MS, C18, 1 mm x 150 mm, 2.0 μm; Thermo Fisher Scientific) connected to an Vanquish Neo UHPLC system and a Q Exactive Plus Orbitrap mass spectrometer (Thermo Fisher Scientific) equipped with a heated electrospray ionisation (HESI) probe. The elution was performed with a gradient of 20 mins. During 0–1 minutes, elution starts with 30% B and increases to 100% in a linear gradient over 14 mins. 100% B is maintained for 5 mins. The flow rate was set to 40 µl/min. MS spectra of lipids were acquired in full-scan/data-dependent MS2 mode. The maximum injection time for full scans was 100 ms, with a target value of 3,000,000 at a resolution of 70,000 and a mass range of 200–1400 m/z in both positive and negative modes. The 10 most intense ions from the survey scan were selected and fragmented with HCD with a stepped collision energy of 25, 30 and 35. Target values for MS/MS were set at 100,000 with a maximum injection time of 50 ms at a resolution of 35,000. To avoid repetitive sequencing, the dynamic exclusion of sequenced lipids was set at 10 s. Peaks were analyzed using the Lipid Search algorithm (MKI, Tokyo, Japan). Peaks were defined through raw files, product ion and precursor ion accurate masses. Candidate molecular species were identified by database (>1,500,000 entries) search of positive (+H^+^; +NH_4_^+^) or negative ion adducts (-H^-^;+COOH^-^). Mass tolerance was set to five ppm for the precursor mass. Samples were aligned within a 0.5 min time window and the results combined in a single report. Lipid standards were used for the calculation of lipid concentrations (Equisplash, Avanti Polar Lipids). Internal standard was used for the normalization between samples. From the intensities of lipid standards and lipid classes, absolute values for each lipid in pmol/µg protein were calculated. All LC-MS/MS lipidomic raw data are provided in the Data Table.

### Targeted analysis of LCBs and ceramides by LC-MS/MS

The targeted analysis of LCBs and ceramides by LC-MS/MS was performed as previously described (Esch et al., 2020, Esch et al., 2023). An external standard curve was prepared using dihydrosphingosine 18:0 (DHS; Avanti Polar Lipids), phytosphingosine 18:0 (PHS; Avanti Polar Lipids) and ceramide t18:0/24:0 (Avanti Polar Lipids/Cayman). Samples were analyzed on a QTRAP 5500 LC-MS/MS (SCIEX) mass spectrometer connected to a Shimadzu Nexera HPLC system and an Accucore C30 LC column (150 mm × 2.1 mm 2.6µm Solid Core; Thermo Fisher Scientific) in positive mode. For the gradient, 40% B for 0.1min was used followed by an increase from 40% to 50% over 1.4 min. Afterwards, buffer B was increased from 50% to 100% over 1.5 min. 100% B was used for 1min and decreased to 40% B for 0.1 min. Then, 40% B was kept until the end of the gradient. A constant flow rate of 0.4 ml/min was used with a total analysis time of 6 min and an injection volume of 2 μl. The MS data were measured in positive ion, scheduled MRM mode without detection windows. The SciexOS software was used for evaluation. The internal standard was used for normalization. Lipid standards were used for the calculation of lipid concentrations. From the intensities of lipid standards, absolute values for each lipid in pmol/µg protein were calculated. For the metabolic flux analysis, measured OD_600_ units were used for correction of the used cell number, resulting in final pmol/OD_600_ units. All LC-MS/MS lipidomic raw data are provided in the DataTable.

### Analysis of ^3^H-myo-inositol-labelled lipids

^3^H-labelled phosphatidylinositol and complex sphingolipid metabolism was analysed by established procedures, as previously described (Gaynor et al., 1999). Briefly, 5 OD_600_ cell equivalents grown to mid-log phase in YND media were washed twice in YND media lacking inositol and pre-incubated for 15 min prior to labelling with 50 μCi of *myo*-[2-^3^H]-inositol for 15 min. Cells were then chased by the addition of 50 μg/ml myo-inositol and metabolically inhibited after a 45 min by the addition of NaN_3_ and NaF (10 mM final concentration each) and placed on ice. Cells were then collected by centrifugation at 4°C, washed in 10 mM NaN_3_/NaF, and resuspended in 500 μl of CHCl_3_/CH_3_OH/H_2_O, 10:10:3 (vol/vol) extraction solvent. Cell suspensions were mechanically lysed by adding glass beads and subsequent rounds of vortexing (four times for 30 sec each). Lysates were centrifuged at 13,000 x *g* for 5 min, and the organic phase was collected and transferred to a new tube. An additional 300 μl of extraction solvent was added to the remaining aqueous phase/glass beads, vortexed 30 sec, and centrifuged as before. The secondary resulting organic phase was collected and pooled with the first organic phase. The pooled lipids were dried in a speed-vac at room temperature. For desalting, the dried lipids were resuspended in 150 μl H_2_O-saturated butanol and extracted with 75 μl H_2_O. The organic phase was collected, and the aqueous phase was re-extracted with an additional 75 μl H_2_O-saturated butanol. The pooled organic phases were dried in a speed-vac and resuspended in 50 μl extraction solvent and aliquots were used to determine total [^3^H]-inositol incorporation by scintillation counting. Equivalent samples were applied to Whatman Linear K6D silica gel thin-layer chromatography (TLC) plates and resolved in CHCl_3_/CH_3_OH/0.25% KCl, 55:45:10 [vol/vol]. Radioactive bands were visualized by X-ray film after treatment with En3Hance (NEN Life Science Products). Phosphatidylinositol, lyso-phosphatidylinositol, and complex sphingolipid levels were determined by densitometry using Fiji (measurement function) (Schindelin et al., 2012). Raw data measurements are provided in the Data Table.

### Western blot analysis of Gas1p

Preparation of cell lysates and western blot analysis were performed as described previously (Kajiwara et al., 2014, Ikeda et al., 2020). Blots were probed with rabbit polyclonal antibodies against Gas1p and a peroxidase-conjugated affinity-purified anti-rabbit IgG antibody.

### Live cell imaging

Live yeast cell imaging experiments were performed on mid-log phase yeast cultures in synthetic media (YND) containing the appropriate nutrient supplements. Imaging data were acquired using a PerkinElmer Ultraview Vox spinning disk confocal microscopy system, equipped with a Nikon TiE inverted stand, a 100x CFI Plan Apochromat VC oil-immersion objective lens (1.4 NA), a Yokogawa CSU-X1 spinning disk scan head, a Hamamatsu C9100-13 EMCCD camera, Prior NanoscanZ Piezo Focus System, and the Nikon Perfect Focus System. All images were collected as square images with 512 × 512 pixels.

### Quantitative image analysis

Quantitative image analyses were conducted using Fiji (Schindelin et al., 2012). The number of Sec13-RFP puncta per cell (in mid-plane optical sections) was calculated using the Find Maxima function on FIJI, using appropriate noise tolerance settings on confocal image sections. Fluorescence intensity analyses (GCaMP3, GFP-Orm1, and GFP-Orm2; Figures S5 and S6) were performed by generating regions of interest around cells (F_total cell_) based upon brightfield images. Background fluorescence was determined by measuring the fluorescence intensity in equivalent areas without cells (F_background_). The background intensity was subtracted from the total cell intensity to determine the specific fluorescence intensity (F_specific_). Raw data are provided in the Data Table.

### Quantitative flow cytometry

Flow cytometry experiments to quantitatively monitor GCaMP3 fluorescence were performed using a BD Accuri C6 flow cytometer as previously described (Omnus et al., 2016, Thomas et al., 2022). Yeast strains expressing a cytoplasmic GCaMP3 reporter or an empty vector were grown to mid-log phase at 30°C in YND media. Background fluorescence was measured using strains containing the empty vector only. 100,000 cells were analyzed from each sample in at least three independent experiments. Raw data are provided in the Data Table.

## Statistical analyses

Statistical analyses were carried out using GraphPad Prism. To compare the mean of two groups (Figures 1C, 4B, S4, and 5D), a two-tailed unpaired *t*-test was used. To compare the mean of multiple groups, we used one-way ANOVA followed by Dunnett’s test (Figures 1B, S1, 3, 4D, 5B, S5, 6, and S6) or Tukey–Kramer multiple comparisons (Figures 2, S2, and 6).

## Acknowledgments

We thank Scott Emr, Ruth Collins, Martha Cyert, and Peter Walter for generously providing reagents. We are also grateful to Jeremy Thorner and Nozomu Kono for providing helpful comments on the manuscript.

Christopher J. Stefan is supported by UKRI Biotechnology and Biological Sciences Research Council funding, award reference BB/X017184/1. Kouchi Funato is supported by Grants-in-Aid for Scientific Research from the Japan Society for the Promotion of Science, Japan (21K19088). Florian Fröhlich is supported by the Heisenberg program of the German research foundation (DFG; project number 491484150). This work was also supported by the CRC 1557 project Z1 (project number 516911785 to F.F.).

## Supplemental Figure Legends

**Figure S1.**
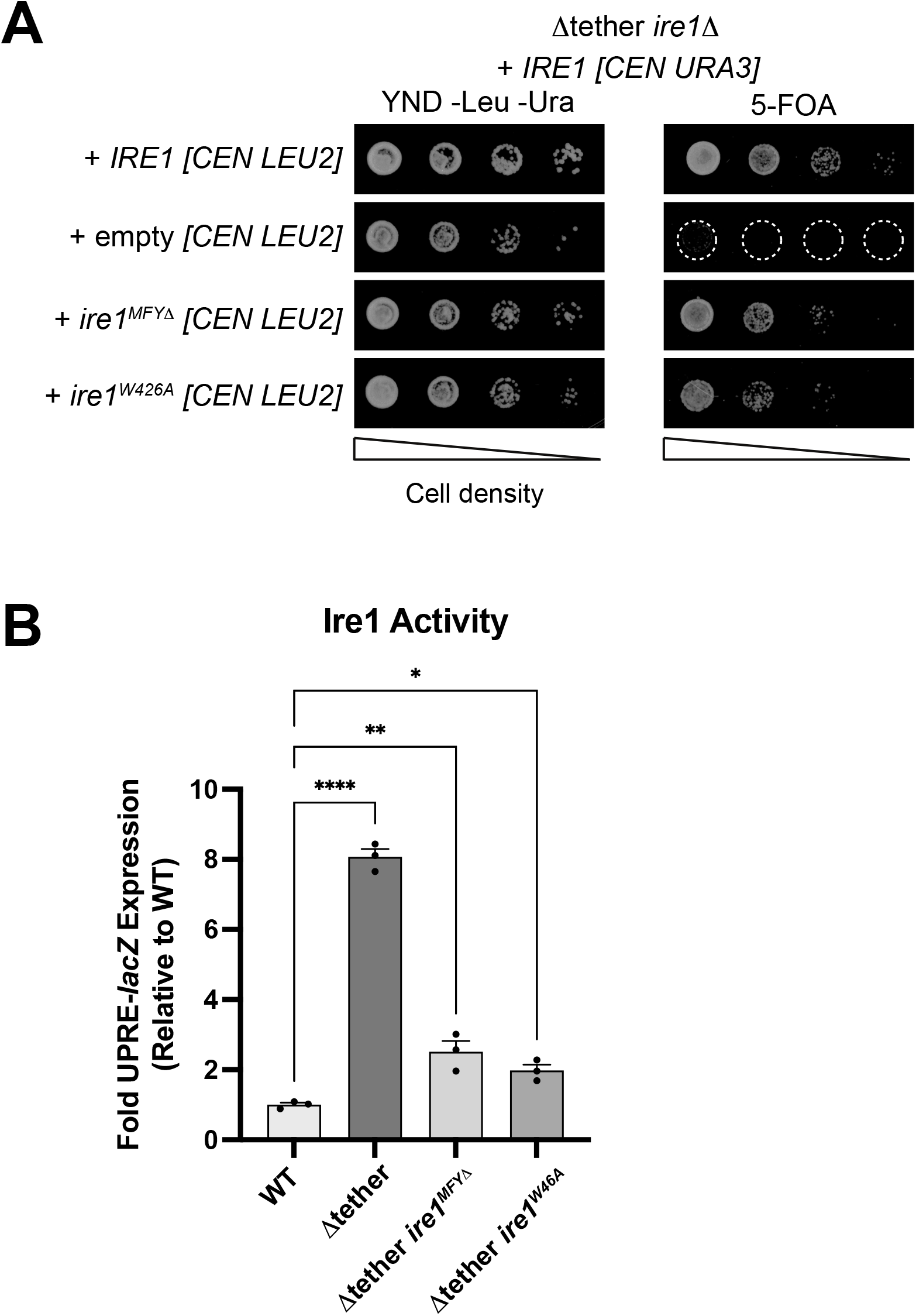
Moderate Ire1 activity in response to membrane lipid stress is sufficient for cell growth upon loss of ER-PM contacts. (A) The Δtether *ire1*Δ cells co-transformed with an *URA3*-marked plasmid expressing wild-type Ire1 and either an empty *LEU2*-marked *CEN* vector or *LEU2*-marked *CEN* vector expressing wild-type Ire1, Ire1^MFYΔ^, or Ire1^W426A^ were spotted (in 10-fold serial dilutions) onto agar media containing 5-FOA to select against the *URA3*-marked *IRE1* plasmid. The Δtether *ire1*Δ cells exhibited a severe growth defect after 2 days. Expression of either Ire1^MFYΔ^ or Ire1^W426A^, which are impaired in UPR, was sufficient to rescue the impaired growth of the Δtether *ire1*Δ cells. (B) Induction of the *UPRE-lacZ* reporter was measured by β-galactosidase activity in the indicated strains. Expression of Ire1^MFYΔ^ or Ire1^W426A^ induce modest activation (2-fold above wild-type basal levels) of the *UPRE-lacZ* reporter in Δtether *ire1*Δ cells. Data represents mean +/-standard error (N = 3). ****, p < 0.0001; **, p < 0.0021; *, p < 0.0332.

**Figure S2.**
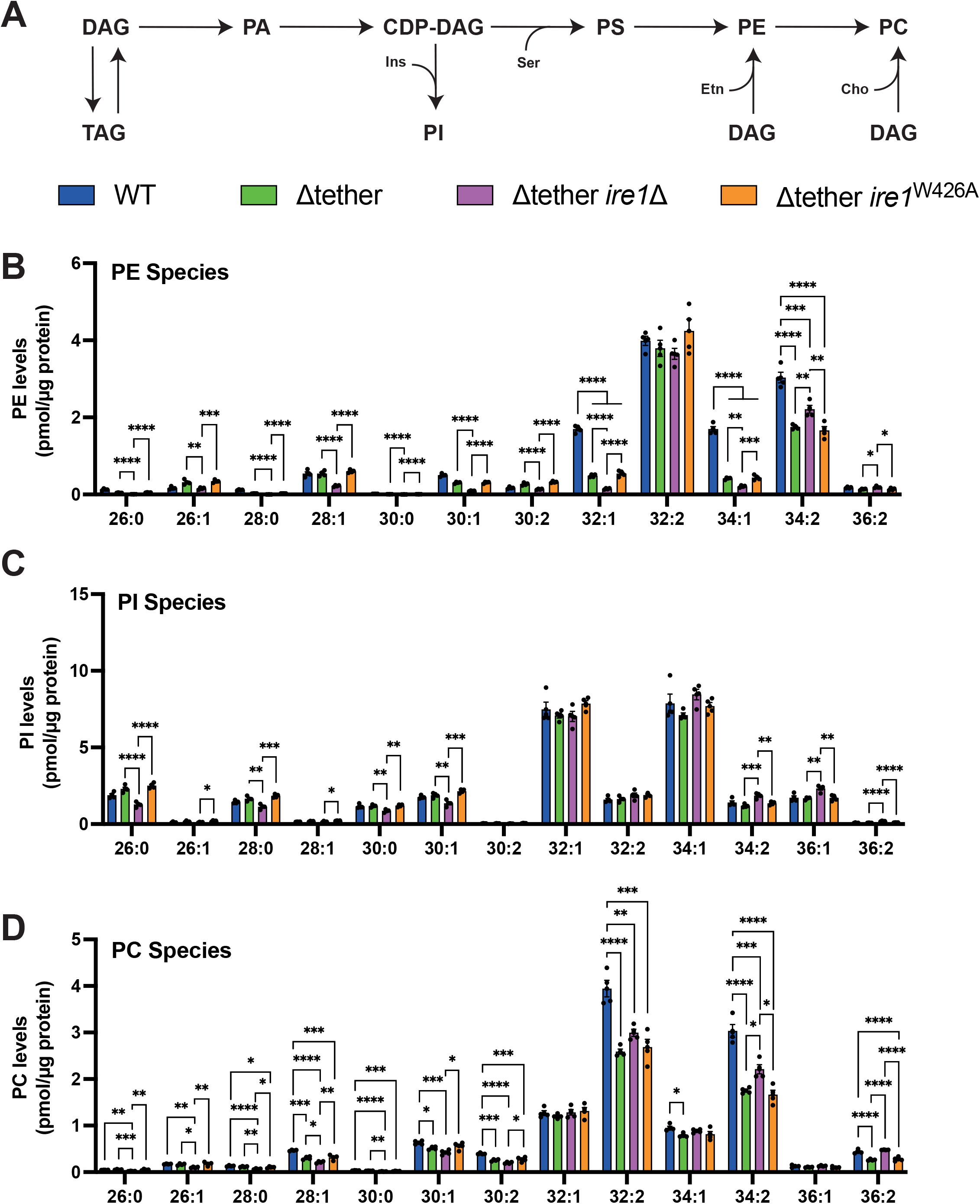
Ire1-mediated membrane stress responses (MSR) sustain pools of saturated and mono-saturated glycerophospholipids upon loss of ER-PM contacts. (A) Schematic of major glycerophospholipid (GPL) biosynthesis pathways in budding yeast. (B-D) Species-level lipidomics analysis (total acyl chain length:double bond number) of PE species (B), PI species (C), and PC species (D) in wild-type (WT), Δtether, Δtether *ire1*Δ and Δtether *ire1*^W426A^ cells. Data represents mean +/-standard error (N=4 independent experiments). ****, p > 0.0001; ***, p > 0.0002; **, p > 0.0021; *, p < 0.0332; ns, not significant. Abbreviations: CDP, cytidine diphosphate; DAG, diacylglycerol; PA, phosphatidic acid; PC, phosphatidylcholine; PE, phosphatidylethanolamine; PI, phosphatidylinositol; PS, phosphatidylserine; TAG, triacylglycerol

**Figure S3.**
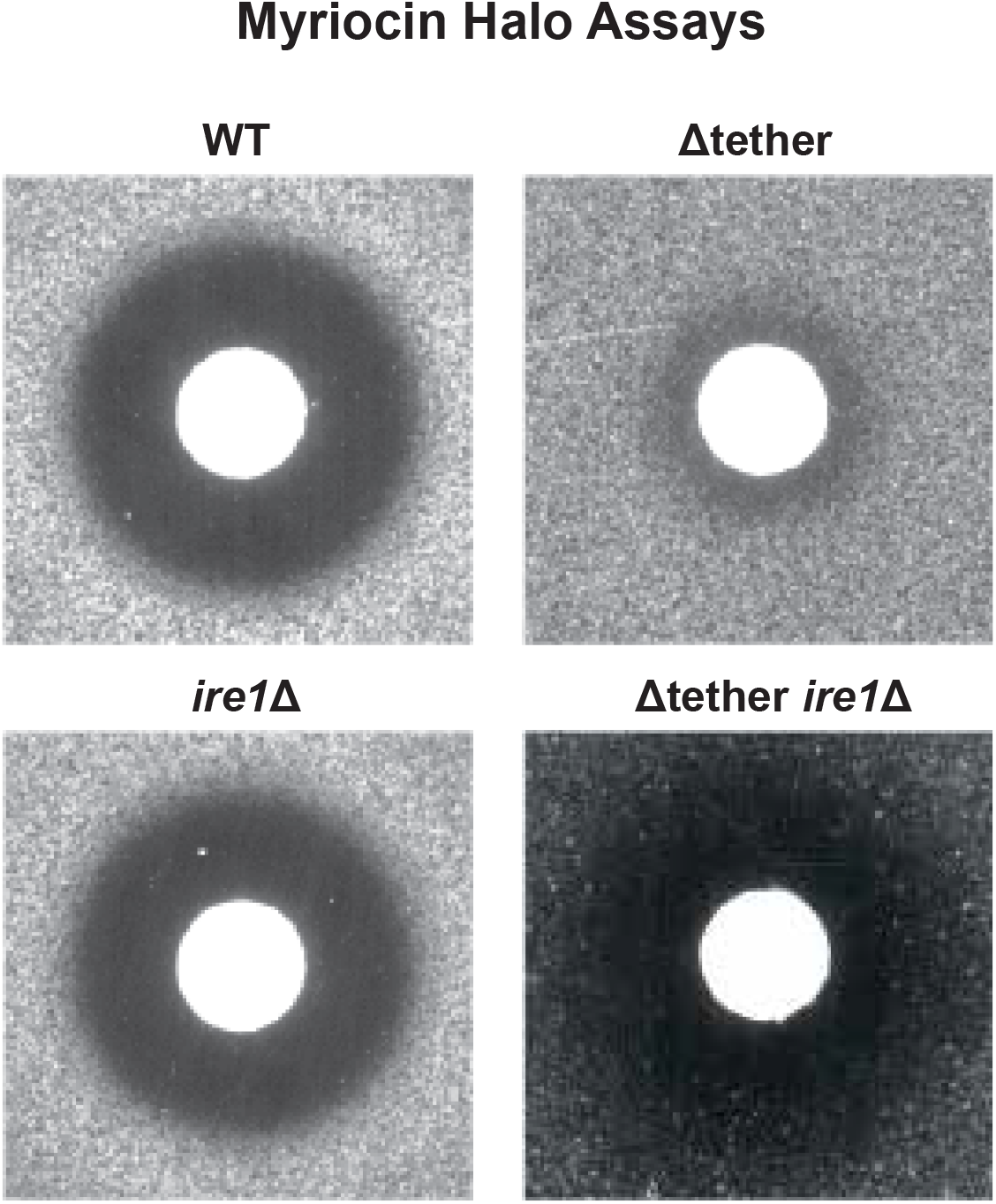
The Δtether cells display myriocin resistance in an Ire1-dependent fashion. Wild-type (WT), Δtether, Δtether *ire1*Δ, and *ire1*Δ cells were plated as lawns in agar and then overlaid with sterile filter discs containing myriocin, a potent SPT inhibitor (3 μl of a 1 mg/ml stock of myriocin in DMSO). After incubation at 30°C for 3 days, the plates were imaged. The Δtether cells show resistance to myriocin, as compared to WT, *ire1*Δ, and Δtether *ire1*Δ cells that form larger zones of growth inhibition (halos).

**Figure S4.**
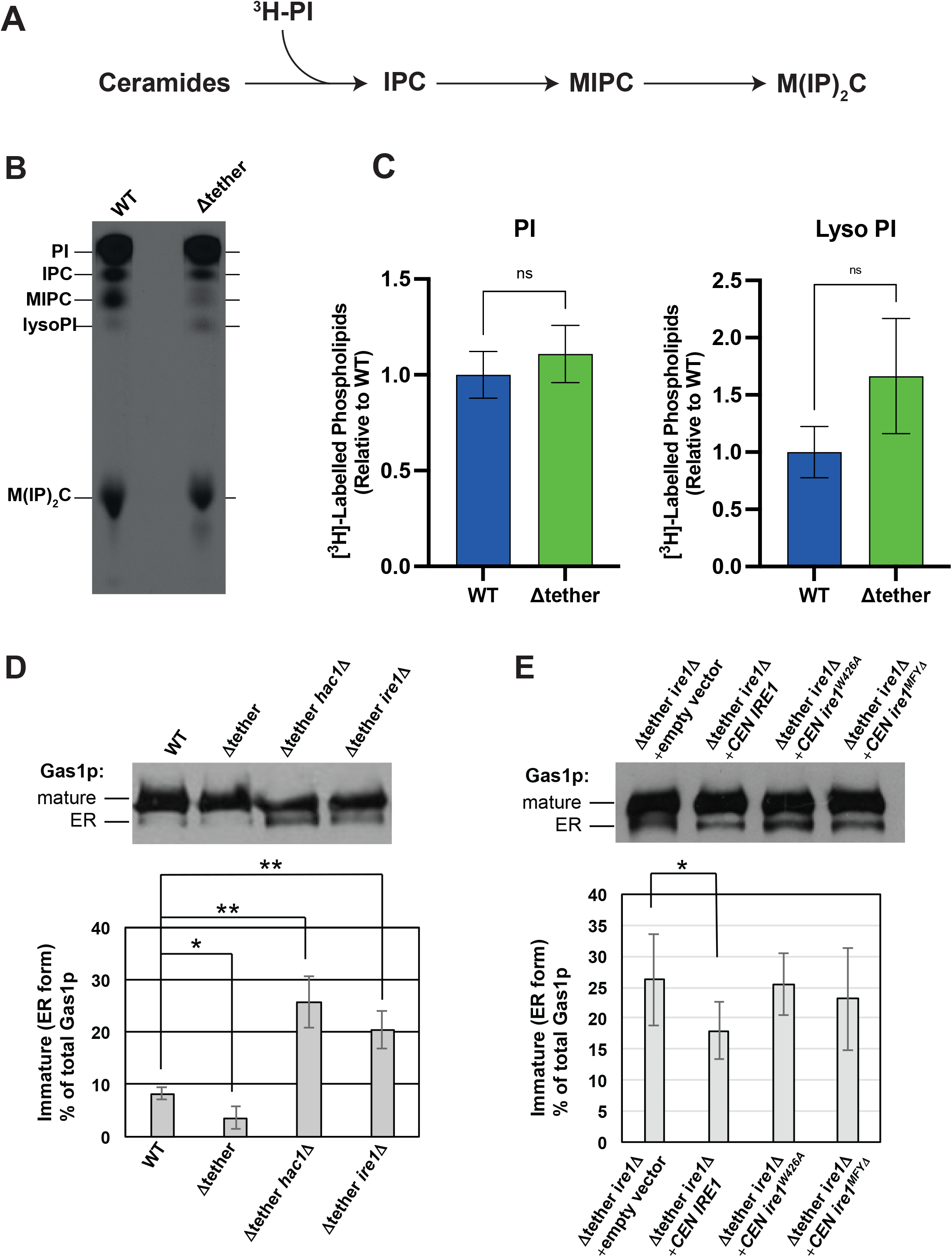
Ire1 MSR does not upregulate ER to Golgi transport in the Δtether cells. (A) Schematic displaying incorporation of ^3^H-myo-inositol into phosphatidylinositol and complex sphingolipids. (B) Representative TLC analysis of ^3^H inositol-labelled of lipids in wild-type (WT) and Δtether cells. Abbreviations: IPC, inositolphosphoceramides; MIPC and M(IP)_2_C, mannosylinositolphosphoceramides; Lyso PI, lyso-phosphatidylinositol; PI, phosphatidylinositol. (C) Quantitation of ^3^H-labelled PI and ^3^H-labelled Lyso-PI in wild-type (WT) and Δtether cells. Data represent mean +/-standard error (N=4). ns, not significant. (D) The Ire1 and Hac1 proteins control steady-state levels of immature forms of the Gas1 protein (Gas1p) in the ER. (E) Ire1-dependent UPR (impaired in the Ire1^W426A^ and Ire1^MFYΔ^ mutant proteins), but not Ire1-mediated MSR (retained by the Ire1^W426A^ and Ire1^MFYΔ^ mutant proteins), is required for control of immature Gas1p in the ER. (D and E) Cell lysates from the indicated strains were prepared, subjected to SDS-PAGE, and analyzed by western blotting using antibodies against Gas1p. Mature, mature forms of Gas1p generated in Golgi compartments; ER, immature ER form of Gas1p. *p < 0.05, **p < 0.01, ***p < 0.001.

**Figure S5.**
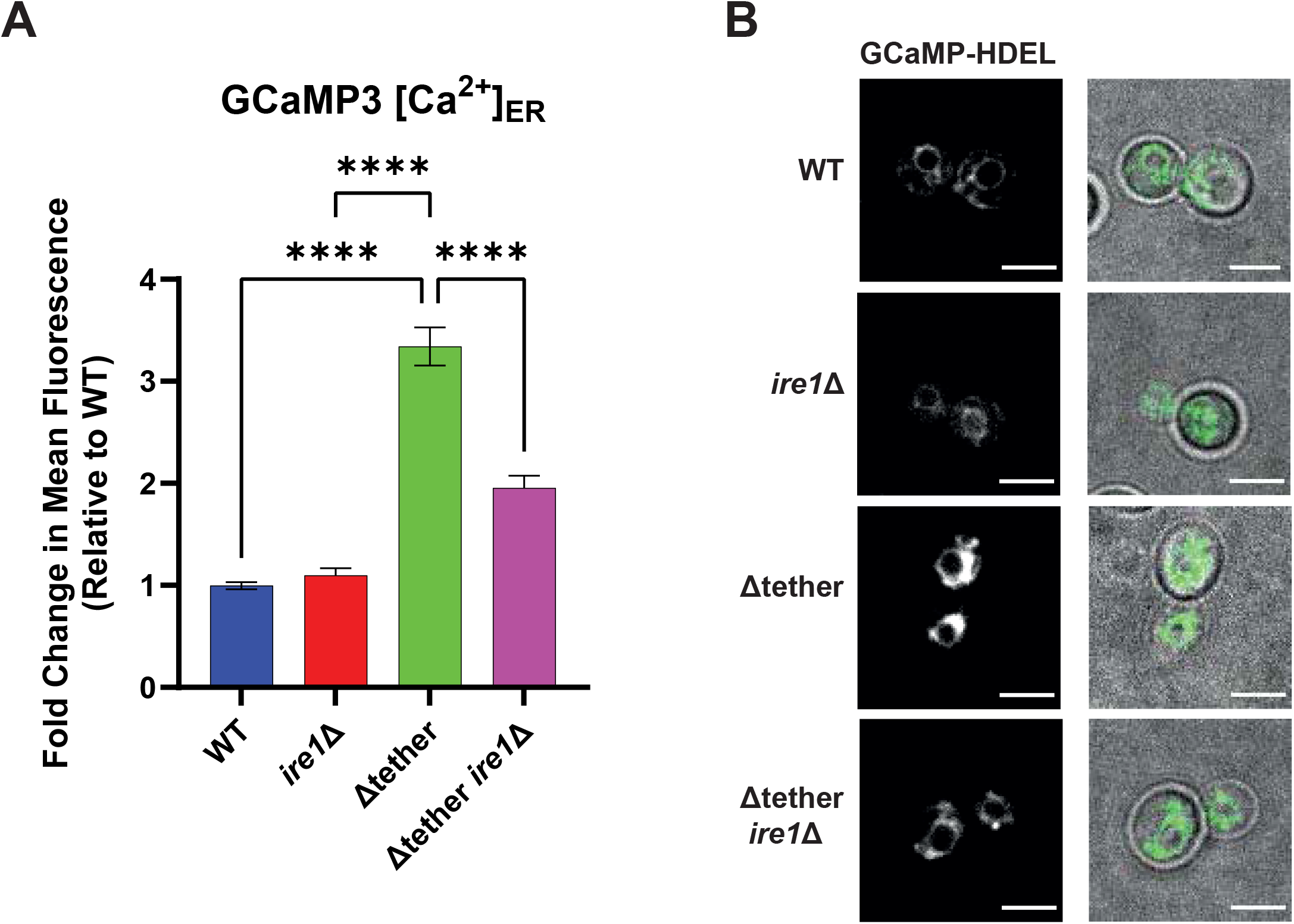
Ire1 regulates Ca^2+^ dynamics in the Δtether cells. (A) Quantification of ER-targeted GCaMP3-HDEL Ca^2+^ reporter specific intensities in wild-type (WT), Δtether, Δtether *ire1*Δ, and *ire1*Δ cells. (B) Representative images of the ER-targeted GCaMP3-HDEL Ca^2+^ reporter in WT, Δtether, Δtether *ire1*Δ and *ire1*Δ cells. Scale bar, 4μm. Data represent mean +/-standard error (N=4). Total number of cells analysed: WT n=208, Δtether n=268, Δtether *ire1*Δ n=115, *ire1*Δ n=108. ns, not significant; ****, p < 0.0001.

**Figure S6.**
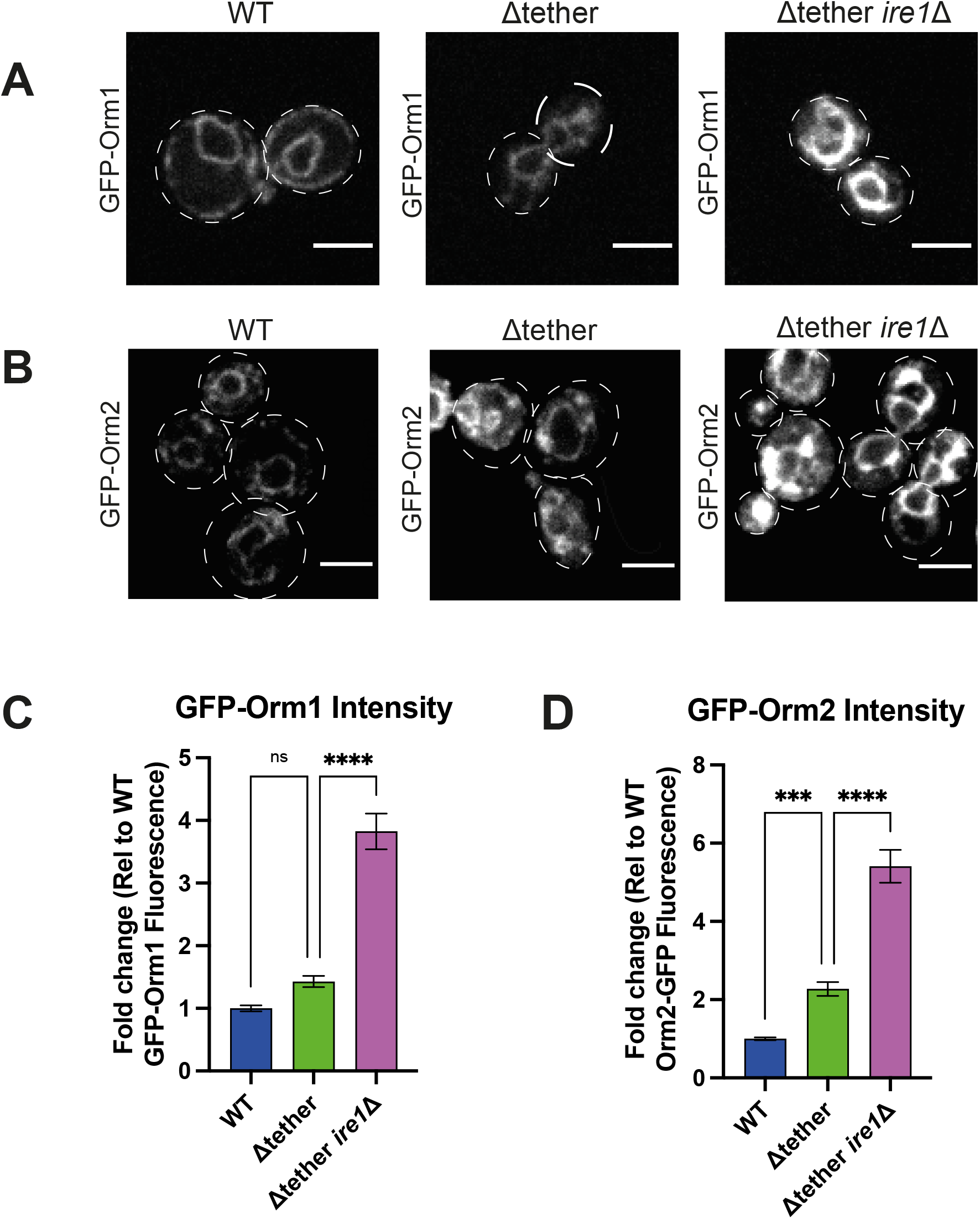
Ire1 modulates Orm1 and Orm2 protein levels in the Δtether cells. Quantitative analyses of GFP-Orm1 (A, C) and GFP-Orm2 (B, D) steady-state levels in wild-type (WT), Δtether, and Δtether *ire1*Δ cells, as determined by confocal microscopy. Data represent mean +/-standard error (N=3). Total number of cells analysed: WT n=197, Δtether n=189, Δtether *ire1*Δ n=199. ****, p < 0.0001; ***, p < 0.0002; ns, not significant. Scale bar, 4 μm.

## References

Aragon, T., Van Anken, E., Pincus, D., Serafimova, I. M., Korennykh, A. V., Rubio, C. A. & Walter, P. 2009. Messenger RNA targeting to endoplasmic reticulum stress signalling sites. Nature, 457, 736–40.

Ariyama, H., Kono, N., Matsuda, S., Inoue, T. & Arai, H. 2010. Decrease in membrane phospholipid unsaturation induces unfolded protein response. J Biol Chem, 285, 22027–35

Aronova, S., Wedaman, K., Aronov, P. A., Fontes, K., Ramos, K., Hammock, B. D. & Powers, T. 2008. Regulation of ceramide biosynthesis by Tor complex 2. Cell Metab, 7, 148–58.

Bao, X., Koorengevel, M. C., Groot Koerkamp, M. J. A., Homavar, A., Weijn, A., Crielaard, S., Renne, M. F., Lorent, J. H., Geerts, W. J., Surma, M. A., Mari, M., Holstege, F. C. P., Klose, C. & De Kroon, A. 2021. Shortening of membrane lipid acyl chains compensates for phosphatidylcholine deficiency in choline-auxotroph yeast. Embo J, 40, e107966.

Barisch, C., Holthuis, J. C. M. & Cosentino, K. 2023. Membrane damage and repair: a thin line between life and death. Biol Chem, 404, 467–490.

Bonilla, M., Nastase, K. K. & Cunningham, K. W. 2002. Essential role of calcineurin in response to endoplasmic reticulum stress. Embo J, 21, 2343–53.

Borgese, N., Iacomino, N., Colombo, S. F. & Navone, F. 2021. The Link between Vapb Loss of Function and Amyotrophic Lateral Sclerosis. Cells, 10.

Breslow, D. K., Collins, S. R., Bodenmiller, B., Aebersold, R., Simons, K., Shevchenko, A., Ejsing, C. S. & Weissman, J. S. 2010. Orm family proteins mediate sphingolipid homeostasis. Nature, 463, 1048–53.

Carreras-Sureda, A., Zhang, X., Laubry, L., Brunetti, J., Koenig, S., Wang, X., Castelbou, C., Hetz, C., Liu, Y., Frieden, M. & Demaurex, N. 2023. The Er stress sensor IRE1 interacts with STIM1 to promote store-operated calcium entry, T cell activation, and muscular differentiation. Cell Rep, 42, 113540.

Chen, H. J., Anagnostou, G., Chai, A., Withers, J., Morris, A., Adhikaree, J., Pennetta, G. & De Belleroche, J. S. 2010. Characterization of the properties of a novel mutation in Vapb in familial amyotrophic lateral sclerosis. J Biol Chem, 285, 40266–81.

Collado, J., Kalemanov, M., Campelo, F., Bourgoint, C., Thomas, F., Loewith, R., Martinez-Sanchez, A., Baumeister, W., Stefan, C. J. & Fernandez-Busnadiego, R. 2019. Tricalbin-Mediated Contact Sites Control Er Curvature to Maintain Plasma Membrane Integrity. Dev Cell, 51, 476–487 e7.

Contreras, F. X., Villar, A. V., Alonso, A. & Goni, F. M. 2009. Ceramide-induced transbilayer (flip-flop) lipid movement in membranes. Methods Mol Biol, 462, 155–65.

Cox, J. S. & Walter, P. 1996. A novel mechanism for regulating activity of a transcription factor that controls the unfolded protein response. Cell, 87, 391–404.

Cunningham, K. W. & Fink, G. R. 1994. Calcineurin-dependent growth control in Saccharomyces cerevisiae mutants lacking PMC1, a homolog of plasma membrane Ca2+ ATPases. J Cell Biol, 124, 351–63.

D’ambrosio, J. M., Albanese, V., Lipp, N. F., Fleuriot, L., Debayle, D., Drin, G. & Copic, A. 2020. Osh6 requires Ist2 for localization to Er-Pm contacts and efficient phosphatidylserine transport in budding yeast. J Cell Sci, 133.

Ejsing, C. S., Sampaio, J. L., Surendranath, V., Duchoslav, E., Ekroos, K., Klemm, R. W., Simons, K. & Shevchenko, A. 2009. Global analysis of the yeast lipidome by quantitative shotgun mass spectrometry. Proc Natl Acad Sci U S A, 106, 2136–41.

Epstein, S., Kirkpatrick, C. L., Castillon, G. A., Muniz, M., Riezman, I., David, F. P. A., Wollheim, C. B. & Riezman, H. 2012. Activation of the unfolded protein response pathway causes ceramide accumulation in yeast and Ins-1e insulinoma cells. J Lipid Res, 53, 412–420.

Ernst, R., Renne, M. F., Jain, A. & Von Der Malsburg, A. 2024. Endoplasmic Reticulum Membrane Homeostasis and the Unfolded Protein Response. Cold Spring Harb Perspect Biol, 16.

Esch, B. M., Limar, S., Bogdanowski, A., Gournas, C., More, T., Sundag, C., Walter, S., Heinisch, J. J., Ejsing, C. S., Andre, B. & Frohlich, F. 2020. Uptake of exogenous serine is important to maintain sphingolipid homeostasis in Saccharomyces cerevisiae. PLos Genet, 16, e1008745.

Esch, B. M., Walter, S., Schmidt, O. & Frohlich, F. 2023. Identification of distinct active pools of yeast serine palmitoyltransferase in sub-compartments of the Er. J Cell Sci, 136.

Filosto, S., Fry, W., Knowlton, A. A. & Goldkorn, T. 2010. Neutral sphingomyelinase 2 (nSMase2) is a phosphoprotein regulated by calcineurin (PP2b). J Biol Chem, 285, 10213–22.

Fu, S., Yang, L., Li, P., Hofmann, O., Dicker, L., Hide, W., Lin, X., Watkins, S. M., Ivanov, A. R. & Hotamisligil, G. S. 2011. Aberrant lipid metabolism disrupts calcium homeostasis causing liver endoplasmic reticulum stress in obesity. Nature, 473, 528–31.

Fun, X. H. & Thibault, G. 2020. Lipid bilayer stress and proteotoxic stress-induced unfolded protein response deploy divergent transcriptional and non-transcriptional programmes. Biochim Biophys Acta Mol Cell Biol Lipids, 1865, 158449.

Funato, K., Lombardi, R., Vallee, B. & Riezman, H. 2003. Lcb4p is a key regulator of ceramide synthesis from exogenous long chain sphingoid base in Saccharomyces cerevisiae. J Biol Chem, 278, 7325–34.

Gardner, B. M. & Walter, P. 2011. Unfolded proteins are Ire1-activating ligands that directly induce the unfolded protein response. Science, 333, 1891–4.

Gaynor, E. C., Mondesert, G., Grimme, S. J., Reed, S. I., Orlean, P. & Emr, S. D. 1999. MCD4 encodes a conserved endoplasmic reticulum membrane protein essential for glycosylphosphatidylinositol anchor synthesis in yeast. Mol Biol Cell, 10, 627–48.

Gkogkas, C., Middleton, S., Kremer, A. M., Wardrope, C., Hannah, M., Gillingwater, T. H. & Skehel, P. 2008. Vapb interacts with and modulates the activity of ATF6. Hum Mol Genet, 17, 1517–26.

Greenberg, M. L., Klig, L. S., Letts, V. A., Loewy, B. S. & Henry, S. A. 1983. Yeast mutant defective in phosphatidylcholine synthesis. J Bacteriol, 153, 791–9.

Gururaj, C., Federman, R. S. & Chang, A. 2013. Orm proteins integrate multiple signals to maintain sphingolipid homeostasis. J Biol Chem, 288, 20453–63.

Gustavsson, M., Traaseth, N. J. & Veglia, G. 2011. Activating and deactivating roles of lipid bilayers on the Ca(2+)-ATPase/phospholamban complex. Biochemistry, 50, 10367–74.

Haak, D., Gable, K., Beeler, T. & Dunn, T. 1997. Hydroxylation of Saccharomyces cerevisiae ceramides requires Sur2p and Scs7p. J Biol Chem, 272, 29704–10.

Halbleib, K., Pesek, K., Covino, R., Hofbauer, H. F., Wunnicke, D., Hanelt, I., Hummer, G. & Ernst, R. 2017. Activation of the Unfolded Protein Response by Lipid Bilayer Stress. Mol Cell, 67, 673–684 e8.

Han, S., Lone, M. A., Schneiter, R. & Chang, A. 2010. Orm1 and Orm2 are conserved endoplasmic reticulum membrane proteins regulating lipid homeostasis and protein quality control. Proc Natl Acad Sci U S A, 107, 5851–6.

Hanaoka, K., Nishikawa, K., Ikeda, A., Schlarmann, P., Sasaki, S., Fujii, S., Yamashita, S., Nakaji, A. & Funato, K. 2024. Membrane contact sites regulate vacuolar fission via sphingolipid metabolism. Elife, 12.

Henry, S. A., Kohlwein, S. D. & Carman, G. M. 2012. Metabolism and regulation of glycerolipids in the yeast Saccharomyces cerevisiae. Genetics, 190, 317–49.

Ho, N., Yap, W. S., Xu, J., Wu, H., Koh, J. H., Goh, W. W. B., George, B., Chong, S. C., Taubert, S. & Thibault, G. 2020. Stress sensor Ire1 deploys a divergent transcriptional program in response to lipid bilayer stress. J Cell Biol, 219.

Hoffmann, P. C., Bharat, T. A. M., Wozny, M. R., Boulanger, J., Miller, E. A. & Kukulski, W. 2019. Tricalbins Contribute to Cellular Lipid Flux and Form Curved Er-Pm Contacts that Are Bridged by Rod-Shaped Structures. Dev Cell, 51, 488–502 e8.

Ikeda, A., Schlarmann, P., Kurokawa, K., Nakano, A., Riezman, H. & Funato, K. 2020. Tricalbins Are Required for Non-vesicular Ceramide Transport at Er-Golgi Contacts and Modulate Lipid Droplet Biogenesis. iScience, 23, 101603.

Ishiwata-Kimata, Y., Promlek, T., Kohno, K. & Kimata, Y. 2013. Bip-bound and nonclustered mode of Ire1 evokes a weak but sustained unfolded protein response. Genes Cells, 18, 288–301.

Jonikas, M. C., Collins, S. R., Denic, V., Oh, E., Quan, E. M., Schmid, V., Weibezahn, J., Schwappach, B., Walter, P., Weissman, J. S. & Schuldiner, M. 2009. Comprehensive characterization of genes required for protein folding in the endoplasmic reticulum. Science, 323, 1693–7.

Jorgensen, J. R., Tei, R., Baskin, J. M., Michel, A. H., Kornmann, B. & Emr, S. D. 2020. Escrt-Iii and Er-Pm contacts maintain lipid homeostasis. Mol Biol Cell, 31, 1302–1313.

Kajiwara, K., Ikeda, A., Aguilera-Romero, A., Castillon, G. A., Kagiwada, S., Hanada, K., Riezman, H., Muniz, M. & Funato, K. 2014. Osh proteins regulate Copii-mediated vesicular transport of ceramide from the endoplasmic reticulum in budding yeast. J Cell Sci, 127, 376–87.

Kanekura, K., Nishimoto, I., Aiso, S. & Matsuoka, M. 2006. Characterization of amyotrophic lateral sclerosis-linked P56s mutation of vesicle-associated membrane protein-associated protein B (Vapb/ALS8). J Biol Chem, 281, 30223–33.

Kato, T., Kubo, A., Nagayama, T., Kume, S., Tanaka, C., Nakayama, Y., Iida, K. & Iida, H. 2017. Genetic analysis of the regulation of the voltage-gated calcium channel homolog Cch1 by the gamma subunit homolog Ecm7 and cortical Er protein Scs2 in yeast. PLos One, 12, e0181436.

Kitai, Y., Ariyama, H., Kono, N., Oikawa, D., Iwawaki, T. & Arai, H. 2013. Membrane lipid saturation activates IRE1alpha without inducing clustering. Genes Cells, 18, 798–809.

Kondo, N., Ohno, Y., Yamagata, M., Obara, T., Seki, N., Kitamura, T., Naganuma, T. & Kihara, A. 2014. Identification of the phytosphingosine metabolic pathway leading to odd-numbered fatty acids. Nat Commun, 5, 5338.

Kono, N., Amin-Wetzel, N. & Ron, D. 2017. Generic membrane-spanning features endow IRE1alpha with responsiveness to membrane aberrancy. Mol Biol Cell, 28, 2318–2332.

Korennykh, A. V., Egea, P. F., Korostelev, A. A., Finer-Moore, J., Zhang, C., Shokat, K. M., Stroud, R. M. & Walter, P. 2009. The unfolded protein response signals through high-order assembly of Ire1. Nature, 457, 687–93.

Körner, C., Schafer, J. H., Esch, B. M., Parey, K., Walter, S., Teis, D., Januliene, D., Schmidt, O., Moeller, A. & Frohlich, F. 2024. The structure of the Orm2-containing serine palmitoyltransferase complex reveals distinct inhibitory potentials of yeast Orm proteins. Cell Rep, 43, 114627.

Kuo, S. C. & Lampen, J. O. 1974. Tunicamycin--an inhibitor of yeast glycoprotein synthesis. Biochem Biophys Res Commun, 58, 287–95.

Landry, C., Costanzo, J. P., Mitne-Neto, M., Zatz, M., Schaffer, A. E., Hatzoglou, M., Muotri, A. R. & Miranda, H. C. 2025. Convergent activation of the integrated stress response and Er-mitochondria uncoupling in Vapb-associated Als. Embo Mol Med, 17, 2299–2331.

Li, Y., Ge, M., Ciani, L., Kuriakose, G., Westover, E. J., Dura, M., Covey, D. F., Freed, J. H., Maxfield, F. R., Lytton, J. & Tabas, I. 2004. Enrichment of endoplasmic reticulum with cholesterol inhibits sarcoplasmic-endoplasmic reticulum calcium ATPase-2b activity in parallel with increased order of membrane lipids: implications for depletion of endoplasmic reticulum calcium stores and apoptosis in cholesterol-loaded macrophages. J Biol Chem, 279, 37030–9.

Li, Y., Liu, D. & Li, S. 2025. IRE1/Xbp1 promotes the clearance of poly(Gr) dipeptide repeats in amyotrophic lateral sclerosis. J Biol Chem, 301, 110764.

Liu, J., Farmer, J. D., Jr., Lane, W. S., Friedman, J., Weissman, I. & Schreiber, S. L. 1991. Calcineurin is a common target of cyclophilin-cyclosporin A and Fkbp-FK506 complexes. Cell, 66, 807–15.

Liu, L. K., Choudhary, V., Toulmay, A. & Prinz, W. A. 2017. An inducible Er-Golgi tether facilitates ceramide transport to alleviate lipotoxicity. J Cell Biol, 216, 131–147.

Loewen, C. J., Young, B. P., Tavassoli, S. & Levine, T. P. 2007. Inheritance of cortical Er in yeast is required for normal septin organization. J Cell Biol, 179, 467–83.

Longtine, M. S., Mckenzie, A., 3rd, Demarini, D. J., Shah, N. G., Wach, A., Brachat, A., Philippsen, P. & Pringle, J. R. 1998. Additional modules for versatile and economical Pcr-based gene deletion and modification in Saccharomyces cerevisiae. Yeast, 14, 953–61.

Lopez-Montero, I., Monroy, F., Velez, M. & Devaux, P. F. 2010. Ceramide: from lateral segregation to mechanical stress. Biochim Biophys Acta, 1798, 1348–56.

Manford, A. G., Stefan, C. J., Yuan, H. L., Macgurn, J. A. & Emr, S. D. 2012. Er-to-plasma membrane tethering proteins regulate cell signaling and Er morphology. Dev Cell, 23, 1129–40.

Matheos, D. P., Kingsbury, T. J., Ahsan, U. S. & Cunningham, K. W. 1997. Tcn1p/Crz1p, a calcineurin-dependent transcription factor that differentially regulates gene expression in Saccharomyces cerevisiae. Genes Dev, 11, 3445–58.

Megyeri, M., Prasad, R., Volpert, G., Sliwa-Gonzalez, A., Haribowo, A. G., Aguilera-Romero, A., Riezman, H., Barral, Y., Futerman, A. H. & Schuldiner, M. 2019. Yeast ceramide synthases, Lag1 and Lac1, have distinct substrate specificity. J Cell Sci, 132.

Megyeri, M., Riezman, H., Schuldiner, M. & Futerman, A. H. 2016. Making Sense of the Yeast Sphingolipid Pathway. J Mol Biol, 428, 4765–4775.

Mori, K., Kawahara, T., Yoshida, H., Yanagi, H. & Yura, T. 1996. Signalling from endoplasmic reticulum to nucleus: transcription factor with a basic-leucine zipper motif is required for the unfolded protein-response pathway. Genes Cells, 1, 803–17.

Mori, K., Sant, A., Kohno, K., Normington, K., Gething, M. J. & Sambrook, J. F. 1992. A 22 bp cis-acting element is necessary and sufficient for the induction of the yeast KAR2 (Bip) gene by unfolded proteins. Embo J, 11, 2583–93.

Muir, A., Ramachandran, S., Roelants, F. M., Timmons, G. & Thorner, J. 2014. TORC2-dependent protein kinase Ypk1 phosphorylates ceramide synthase to stimulate synthesis of complex sphingolipids. Elife, 3.

Nakahara, K., Ohkuni, A., Kitamura, T., Abe, K., Naganuma, T., Ohno, Y., Zoeller, R. A. & Kihara, A. 2012. The Sjogren-Larsson syndrome gene encodes a hexadecenal dehydrogenase of the sphingosine 1-phosphate degradation pathway. Mol Cell, 46, 461–71.

Nenadic, A., Zaman, M. F., Johansen, J., Volpiana, M. W. & Beh, C. T. 2023. Increased Phospholipid Flux Bypasses Overlapping Essential Requirements for the Yeast Sac1p Phosphoinositide Phosphatase and Er-Pm Membrane Contact Sites. J Biol Chem, 299, 105092.

Nishimura, A. L., Al-Chalabi, A. & Zatz, M. 2005. A common founder for amyotrophic lateral sclerosis type 8 (ALS8) in the Brazilian population. Hum Genet, 118, 499–500.

Nishimura, T., Gecht, M., Covino, R., Hummer, G., Surma, M. A., Klose, C., Arai, H., Kono, N. & Stefan, C. J. 2019. Osh Proteins Control Nanoscale Lipid Organization Necessary for Pi(4,5)P(2) Synthesis. Mol Cell, 75, 1043–1057 e8.

Nishimura, T. & Stefan, C. J. 2020. Specialized Er membrane domains for lipid metabolism and transport. Biochim Biophys Acta Mol Cell Biol Lipids, 1865, 158492.

Obara, K. & Kihara, A. 2017. The Rim101 pathway contributes to Er stress adaptation through sensing the state of plasma membrane. Biochem J, 474, 51–63.

Okamura, K., Kimata, Y., Higashio, H., Tsuru, A. & Kohno, K. 2000. Dissociation of Kar2p/Bip from an Er sensory molecule, Ire1p, triggers the unfolded protein response in yeast. Biochem Biophys Res Commun, 279, 445–50.

Omnus, D. J., Manford, A. G., Bader, J. M., Emr, S. D. & Stefan, C. J. 2016. Phosphoinositide kinase signaling controls Er-Pm cross-talk. Mol Biol Cell, 27, 1170–80.

Park, S. W., Zhou, Y., Lee, J., Lee, J. & Ozcan, U. 2010. Sarco(endo)plasmic reticulum Ca2+-ATPase 2b is a major regulator of endoplasmic reticulum stress and glucose homeostasis in obesity. Proc Natl Acad Sci U S A, 107, 19320–5.

Pineau, L., Colas, J., Dupont, S., Beney, L., Fleurat-Lessard, P., Berjeaud, J. M., Berges, T. & Ferreira, T. 2009. Lipid-induced Er stress: synergistic effects of sterols and saturated fatty acids. Traffic, 10, 673–90.

Platzek, A., Odehnalova, K., Schessner, J. P., Borner, G. H. H. & Schuck, S. 2025. Dynamic Organellar Mapping in yeast reveals extensive protein localization changes during Er stress. Nat Commun, 16, 10842.

Prinz, W. A., Toulmay, A. & Balla, T. 2020. The functional universe of membrane contact sites. Nat Rev Mol Cell Biol, 21, 7–24.

Promlek, T., Ishiwata-Kimata, Y., Shido, M., Sakuramoto, M., Kohno, K. & Kimata, Y. 2011. Membrane aberrancy and unfolded proteins activate the endoplasmic reticulum stress sensor Ire1 in different ways. Mol Biol Cell, 22, 3520–32.

Quon, E., Nenadic, A., Zaman, M. F., Johansen, J. & Beh, C. T. 2022. Er-Pm membrane contact site regulation by yeast ORPs and membrane stress pathways. PLos Genet, 18, e1010106.

Quon, E., Sere, Y. Y., Chauhan, N., Johansen, J., Sullivan, D. P., Dittman, J. S., Rice, W. J., Chan, R. B., Di Paolo, G., Beh, C. T. & Menon, A. K. 2018. Endoplasmic reticulum-plasma membrane contact sites integrate sterol and phospholipid regulation. PLos Biol, 16, e2003864.

Rajakumar, T., Hossain, M. A., Stopka, S. A., Micoogullari, Y., Ang, J., Agar, N. Y. R. & Hanna, J. 2024. Dysregulation of ceramide metabolism causes phytoceramide-dependent induction of the unfolded protein response. Mol Biol Cell, 35, ar117.

Renne, M. F. & Ernst, R. 2023. Membrane homeostasis beyond fluidity: control of membrane compressibility. Trends Biochem Sci, 48, 963–977.

Rockenfeller, P. & Gourlay, C. W. 2018. Lipotoxicty in yeast: a focus on plasma membrane signalling and membrane contact sites. Fems Yeast Res, 18.

Saito, S., Ishikawa, T., Ninagawa, S., Okada, T. & Mori, K. 2022. A motor neuron disease-associated mutation produces non-glycosylated Seipin that induces Er stress and apoptosis by inactivating SERCA2b. Elife, 11.

Schäfer, J. H., Korner, C., Esch, B. M., Limar, S., Parey, K., Walter, S., Januliene, D., Moeller, A. & Frohlich, F. 2023. Structure of the ceramide-bound Spots complex. Nat Commun, 14, 6196.

Schindelin, J., Arganda-Carreras, I., Frise, E., Kaynig, V., Longair, M., Pietzsch, T., Preibisch, S., Rueden, C., Saalfeld, S., Schmid, B., Tinevez, J. Y., White, D. J., Hartenstein, V., Eliceiri, K., Tomancak, P. & Cardona, A. 2012. Fiji: an open-source platform for biological-image analysis. Nat Methods, 9, 676-82.

Schmidt, O., Weyer, Y., Baumann, V., Widerin, M. A., Eising, S., Angelova, M., Schleiffer, A., Kremser, L., Lindner, H., Peter, M., Frohlich, F. & Teis, D. 2019. Endosome and Golgi-associated degradation (Egad) of membrane proteins regulates sphingolipid metabolism. Embo J, 38, e101433.

Schneiter, R., Brugger, B., Sandhoff, R., Zellnig, G., Leber, A., Lampl, M., Athenstaedt, K., Hrastnik, C., Eder, S., Daum, G., Paltauf, F., Wieland, F. T. & Kohlwein, S. D. 1999. Electrospray ionization tandem mass spectrometry (Esi-Ms/Ms) analysis of the lipid molecular species composition of yeast subcellular membranes reveals acyl chain-based sorting/remodeling of distinct molecular species en route to the plasma membrane. J Cell Biol, 146, 741–54.

Schuck, S., Prinz, W. A., Thorn, K. S., Voss, C. & Walter, P. 2009. Membrane expansion alleviates endoplasmic reticulum stress independently of the unfolded protein response. J Cell Biol, 187, 525–36.

Shyu, P., Jr., Ng, B. S. H., Ho, N., Chaw, R., Seah, Y. L., Marvalim, C. & Thibault, G. 2019. Membrane phospholipid alteration causes chronic Er stress through early degradation of homeostatic Er-resident proteins. Sci Rep, 9, 8637.

Stathopoulos, A. M. & Cyert, M. S. 1997. Calcineurin acts through the CRZ1/TCN1-encoded transcription factor to regulate gene expression in yeast. Genes Dev, 11, 3432–44.

Stefan, C. P. & Cunningham, K. W. 2013. Kch1 family proteins mediate essential responses to endoplasmic reticulum stresses in the yeasts Saccharomyces cerevisiae and Candida albicans. J Biol Chem, 288, 34861–70.

Surma, M. A., Klose, C., Peng, D., Shales, M., Mrejen, C., Stefanko, A., Braberg, H., Gordon, D. E., Vorkel, D., Ejsing, C. S., Farese, R., Jr., Simons, K., Krogan, N. J. & Ernst, R. 2013. A lipid E-Map identifies Ubx2 as a critical regulator of lipid saturation and lipid bilayer stress. Mol Cell, 51, 519–30.

Suzuki, H., Kanekura, K., Levine, T. P., Kohno, K., Olkkonen, V. M., Aiso, S. & Matsuoka, M. 2009. Als-linked P56s-Vapb, an aggregated loss-of-function mutant of Vapb, predisposes motor neurons to Er stress-related death by inducing aggregation of co-expressed wild-type Vapb. J Neurochem, 108, 973–985.

Thibault, G., Shui, G., Kim, W., Mcalister, G. C., Ismail, N., Gygi, S. P., Wenk, M. R. & Ng, D. T. 2012. The membrane stress response buffers lethal effects of lipid disequilibrium by reprogramming the protein homeostasis network. Mol Cell, 48, 16–27.

Thomas, F. B., Omnus, D. J., Bader, J. M., Chung, G. H., Kono, N. & Stefan, C. J. 2022. Tricalbin proteins regulate plasma membrane phospholipid homeostasis. Life Sci Alliance, 5.

Toulmay, A. & Prinz, W. A. 2012. A conserved membrane-binding domain targets proteins to organelle contact sites. J Cell Sci, 125, 49–58.

Travers, K. J., Patil, C. K., Wodicka, L., Lockhart, D. J., Weissman, J. S. & Walter, P. 2000. Functional and genomic analyses reveal an essential coordination between the unfolded protein response and Er-associated degradation. Cell, 101, 249–58.

Valenzuela, V., Becerra, D., Astorga, J. I., Fuentealba, M., Diaz, G., Bargsted, L., Chacon, C., Martinez, A., Gozalvo, R., Jackson, K., Morales, V., Heras, M. L., Tamburini, G., Petrucelli, L., Sardi, S. P., Plate, L. & Hetz, C. 2025. Artificial enforcement of the unfolded protein response reduces disease features in multiple preclinical models of Als/Ftd. Mol Ther, 33, 1226–1245.

Voeltz, G. K., Sawyer, E. M., Hajnoczky, G. & Prinz, W. A. 2024. Making the connection: How membrane contact sites have changed our view of organelle biology. Cell, 187, 257–270.

Volmer, R. & Ron, D. 2015. Lipid-dependent regulation of the unfolded protein response. Curr Opin Cell Biol, 33, 67–73.

Volmer, R., Van Der Ploeg, K. & Ron, D. 2013. Membrane lipid saturation activates endoplasmic reticulum unfolded protein response transducers through their transmembrane domains. Proc Natl Acad Sci U S A, 110, 4628–33.

Walter, P. & Ron, D. 2011. The unfolded protein response: from stress pathway to homeostatic regulation. Science, 334, 1081–6.

Wolf, W., Kilic, A., Schrul, B., Lorenz, H., Schwappach, B. & Seedorf, M. 2012. Yeast Ist2 recruits the endoplasmic reticulum to the plasma membrane and creates a ribosome-free membrane microcompartment. PLos One, 7, e39703.

Wong, A. K. O., Young, B. P. & Loewen, C. J. R. 2021. Ist2 recruits the lipid transporters Osh6/7 to Er-Pm contacts to maintain phospholipid metabolism. J Cell Biol, 220.

Xie, T., Dong, F., Han, G., Wu, X., Liu, P., Zhang, Z., Zhong, J., Niranjanakumari, S., Gable, K., Gupta, S. D., Liu, W., Harrison, P. J., Campopiano, D. J., Dunn, T. M. & Gong, X. 2024. Collaborative regulation of yeast Spt-Orm2 complex by phosphorylation and ceramide. Cell Rep, 43, 113717.

